# Intratumoral CXCL12 Gradients Contextualize Tumor Cell Invasion, Migration and Immune Suppression in Breast Cancer

**DOI:** 10.1101/2024.10.15.618571

**Authors:** Dimitra P. Anastasiadou, Nicole Couturier, Sakshi Goel, Dimitrios G. Argyris, Stepan Vodopyanov, Luis Rivera-Sanchez, Edgar Gonzalez, Jesse Kreger, Anthony Griffen, Abigail Kazakov, Joseph Burt, Natasha Recoder, Camille L. Duran, Allison S. Harney, Agathe Quesnel, Panagiota S. Filippou, Vasileios P. Lenis, Suryansh Shukla, David Entenberg, Aliona Zintiridou, Xiaoming Chen, Robert J. Eddy, Maja H. Oktay, John S. Condeelis, Nikolaos S. Karagiannis, Andrea Briceno, Hillary Guzik, Rotem Alon, Vera DesMarais, Giorgio Ioannou, Sacha Gnjatic, David. M. Raynolds, Rodney Macedo, Ran Reshef, Hava Gil-Henn, Adam L. MacLean, Evanthia Roussos Torres, Lindsay M. LaFave, Gregoire Lauvau, George S. Karagiannis

**Affiliations:** Department of Microbiology & Immunology, Albert Einstein College of Medicine, Bronx, New York, USA; Tumor Microenvironment & Metastasis Program, Montefiore-Einstein Comprehensive Cancer Center, Bronx, New York, USA; Integrated Imaging Program for Cancer Research, Albert Einstein College of Medicine, Bronx, New York, USA; Department of Cell Biology, Albert Einstein College of Medicine, Bronx, New York, USA; Department of Surgery, Montefiore Medical Center, Bronx, New York, USA; Keck School of Medicine, Department of Medicine, Division of Oncology, Norris Comprehensive Cancer Center, University of Southern California, CA, USA; Department of Quantitative and Computational Biology, University of Southern California, Los Angeles, CA, USA; Department of Pathology, Montefiore Medical Center, Bronx, New York, USA; Gruss-Lipper Biophotonics Center, Albert Einstein College of Medicine, Bronx, New York, USA; Translational Medicine, Cogent Biosciences, Waltham Massachusetts, USA; School of Health & Life Sciences, Teesside University, Middlesbrough, United Kingdom; National Horizons Centre, Teesside University, Darlington, United Kingdom; Laboratory of Biological Chemistry, School of Medicine, Faculty of Health Sciences, Aristotle University of Thessaloniki, Thessaloniki, Greece; Department of Cancer and Genomic Sciences, University of Birmingham, Edgbaston, Birmingham, UK; Centre for Health Data Science, University of Birmingham, Edgbaston, Birmingham, UK; Analytical Imaging Facility, Albert Einstein College of Medicine, Bronx, New York, USA; Gruss Magnetic Resonance Research Center, Albert Einstein College of Medicine, Bronx, New York, USA; Human Immune Monitoring Center (HIMC), Mount Sinai Hospital, New York; Department of Genetics, Albert Einstein College of Medicine, Bronx, NY, USA; Genomics Core, Albert Einstein College of Medicine, Bronx, NY, USA; Columbia Center for Translational Immunology, Department of Medicine, Columbia University Medical Center, New York, New York, USA; The Azrieli Faculty of Medicine, Bar-Ilan University, Safed, Israel; Cancer Dormancy and Tumor Microenvironment Institute, Montefiore-Einstein Comprehensive Cancer Center, Bronx, New York, USA; Marilyn and Stanley M. Katz Institute for Immunotherapy of Cancer and Inflammatory Disorders, Montefiore-Einstein Comprehensive Cancer Center, Bronx, New York, USA; Stem Cell and Cancer Biology Program, Montefiore-Einstein Comprehensive Cancer Center, Bronx, New York, USA; Ruth L. and David S. Gottesman Institute for Stem Cell Research and Regenerative Medicine, Albert Einstein College of Medicine, Bronx, New York, USA

## Abstract

Although the CXCL12/CXCR4 pathway has been prior investigated for its prometastatic and immuno- suppressive roles in the tumor microenvironment, evidence on the spatiotemporal regulation of these hallmarks has been lacking. Here, we demonstrate that CXCL12 forms a gradient specifically around cancer cell intravasation doorways, also known as Tumor Microenvironment of Metastasis (TMEM) doorways, thus facilitating the chemotactic translocation of prometastatic tumor cells expressing CXCR4 toward the perivascular TMEM doorways for subsequent entry into peripheral circulation. Furthermore, we demonstrate that the CXCL12-rich micro-environment around TMEM doorways may create immunosuppressive niches, whereby CD8^+^ T cells, despite being attracted to these regions, often exhibit reduced effector functions, limiting their efficacy. While the CXCL12/CXCR4 pathway can minimally influence the overall composition of immune cell populations, it biases the distribution of CD8^+^ T cells away from TMEM doorways, justifying its prior-established role as immunosuppressive factor for CD8^+^ T cells. Our research suggests that the complex interactions between CXCL12 and the various tumor and immune cell types contributes not only to the completion of the initial steps of the metastatic cascade, but also offers an immunological “sanctuary” to prometastatic tumor cells homed around TMEM doorways. Overall, our study enhances our current understanding on the mechanisms, via which CXCL12 orchestrates tumor cell behavior and immune dynamics, potentially guiding future therapeutic strategies to combat breast cancer metastasis and improve anti-tumor immunity.

## INTRODUCTION

The C-X-C chemokine receptor type 4 (CXCR4) and its ligand, C-X-C motif chemokine 12 (CXCL12) are crucial for embryonic development and adult tissue homeostasis. CXCR4 is expressed by various hematopoietic cells, vascular endothelial cells, endothelial progenitors, and nervous system cells^1–3^. In breast cancer, the CXCL12/CXCR4 pathway regulates tumor cell growth, invasion, and metastasis, and CXCR4 expression correlates with increased tumor progression, grade, and worse survival^4–6^. The CXCR4^+^ tumor cells can migrate towards CXCL12 produced in peripheral organs such as the liver, lungs, brain, and bone marrow, where they can home, survive, and proliferate. Thus, suppressing the CXCL12/CXCR4 axis reduces metastatic burden in mouse models of cancer, suggesting that cancer cells exploit this pathway to metastasize to distant sites^7–12^. However, primary tumors often express high levels of CXCL12, especially under hypoxic conditions, indicating they could also potentially retain CXCR4^+^ cancer cells within their confines. Although the mechanism via which the CXCR4^+^ tumor cells circumvent intratumoral CXCL12 to escape the primary tumor microenvironment is unknown, Balkwill (2004) theorized quite early that microanatomical differences in CXCL12 expression might be sufficient to drive localized invasion and chemotaxis. Such intratumoral CXCL12 niches could theoretically permit CXCR4^+^ tumor cells to navigate through a complex biochemical and molecular tumor microenvironment (TME), allowing them to reach the vasculature for subsequent intravasation. Here, to address this hypothesis, we establish a new paradigm on the involvement of the CXCL12/CXCR4 axis in orchestrating a perivascular niche at the primary tumor microenvironment, uniquely suited to prime the metastatic cascade and provoke an immunosuppressive milieu.

## RESULTS

### CXCL12 and CXCR4 expression in breast carcinoma is tied to the perivascular niche

To define the distribution of CXCL12^+^ and CXCR4^+^ cells in the tumor microenvironment, we conducted single-cell RNA-sequencing (scRNA-seq) in 10-week-old MMTV-PyMT carcinomas, using *MMTV-PyMT; Tie2-GFP* mice **(Figs. 1** **and S1a-c)**. The analysis showed that *Cxcl12* expression was mostly restricted in Tie2^+^ endothelial cells, which otherwise expressed typical endothelial cell markers, such as endomucin (*Emcn*) and *Pecam1/Cd31*, as well as in mesenchymal cells, which expressed Fibronectin (*Fn1*), Platelet-derived growth factor receptors-α and -β (*Pdgfrα/β*), and Fibroblast activation protein (*Fap*), thus consisting of both fibroblasts and pericytes **(Figs. 1a** **and S1d)**. On the other hand, *Cxcr4* expression was observed in distinct myeloid (*Cd68*^+^/*Csf1r*^+^) and lymphoid (*Cd3d*^+^) cell clusters, in *Tie2^+^* endothelial cells, as well as in a few scattered tumor cells within the PyMT^+^ tumor cell cluster, indicating a minority and quite heterogeneous population **(Figs. 1a** **and S1d)**. We partially confirmed these observations in an independent single-cell dataset from an established spontaneously metastatic neu-expressing mouse model, the NT2.5 breast cancer syngeneic model^13^. In this model, *Cxcl12* was notably expressed in both mammary endothelial and cancer-associated fibroblast clusters, while *Cxcr4* expression largely defined the myeloid/lymphoid, endothelial, and tumor cell clusters **(Figs. S2a-c)**.

**Figure 1.**
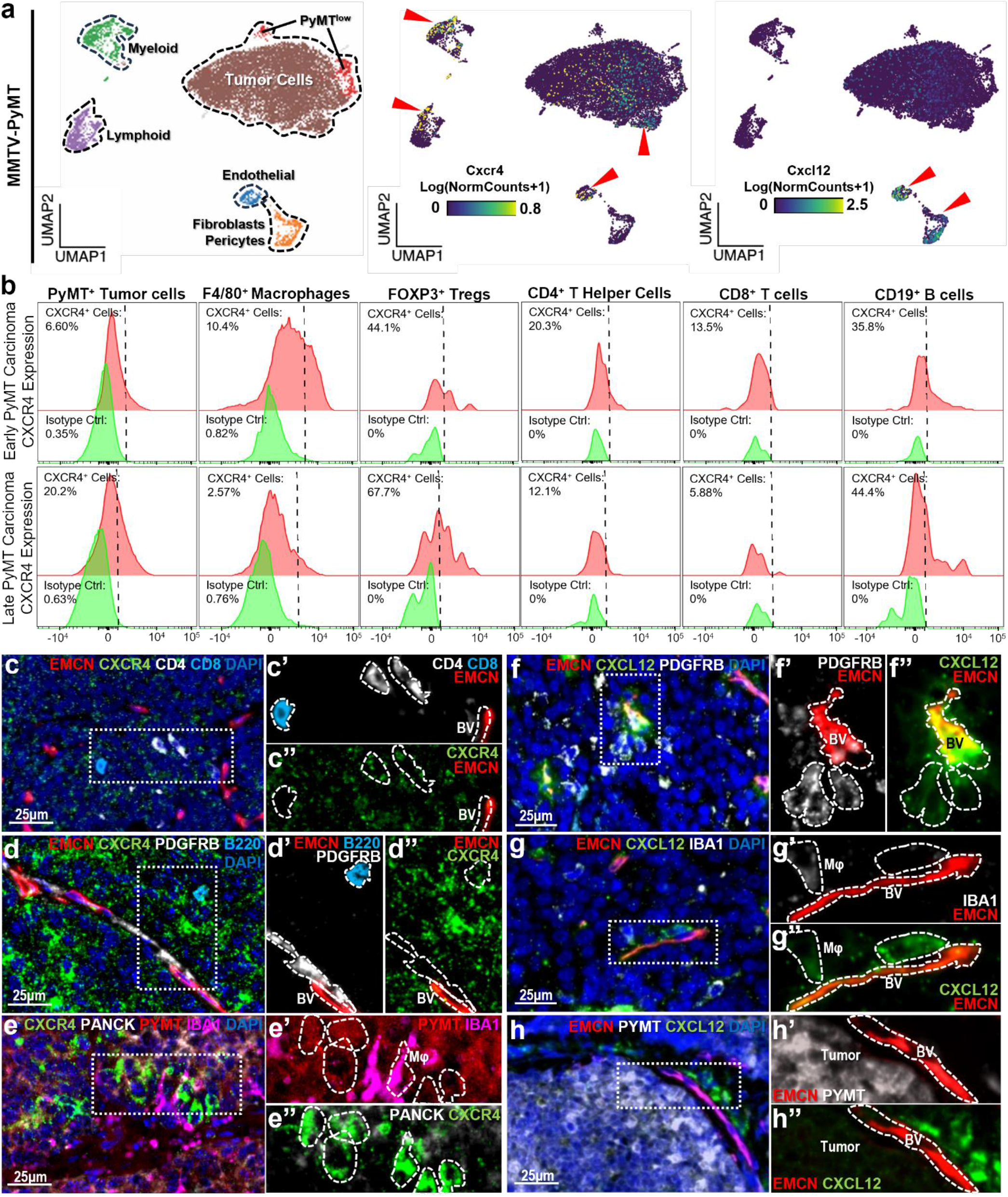
Expression of CXCL12 and CXCR4 in the murine breast tumor microenvironment. **(a)** UMAP visualizations of PyMT carcinomas profiled by scRNA-seq. Left panel: Identification of key cell clusters, using marker genes for tumor cells, endothelial cells, mesenchymal cells (fibroblasts/pericytes), and immune cells (myeloid and lymphoid), as indicated in Supplementary Figure 1. Log normalized expression scores are shown on the UMAP for *Cxcr4* (middle panel) and *Cxcl12* (right panel). Red arrows point to the key clusters and UMAP projections with the highest score for each respective gene. **(b)** Flow cytometry measuring cell-surface CXCR4 expression in gated tumor and immune cell populations at both early-stage (top row) and late-stage (bottom row) MMTV-PyMT carcinomas. The CXCR4^+^ cells are reported as a fraction (%) of the total cells for each cell subset. **(c-h)** Multichannel immunofluorescence using antibodies against a variety of lineage markers, along with CXCR4 (c-e) or CXCL12 (f-h) to track down the cell populations expressing CXCL12 or CXCR4 in MMTV-PyMT carcinomas. Lymphocytes are identified via CD4, CD8 for T cells (panel c) and B220 for B cells (panel d); tumor cells are identified via PANCK or PyMT (panels e, and h); endothelial cells are identified by the expression of EMCN (panels c, d, f, g, and h); mesenchymal cells (pericytes) are identified by the expression of PDGFRB (panels d, f); macrophages are identified by the expression of IBA1 (panels e, and g); all cell nuclei are identified by DAPI expression. Hyphenated and double-hyphenated panels are magnified inserts of the dotted white boxes in the main figure panels. Cells-of-interest in each panel are circumscribed with white dotted lines. BV, blood vessel, Mφ, macrophage.

We extended our observations in MMTV-PyMT mice using high-dimensional flow cytometry against a panel for characterizing tumor immune microenvironments, consisting of both cell lineage and a wide variety of functional markers **(Fig. S3)**. We found that CXCR4 is indeed expressed by a PyMT^+^ tumor cell subset demonstrating slight increase with tumor progression **(Fig. 1b)**, in consistency with previous literature suggesting *de novo* hijacking of the CXCL12/CXCR4 pathway by epithelial tumor cells^14^. As expected, CXCR4 is highly expressed by various immune cell subsets, including F4/80^+^ macrophages (∼10% at early-stage and ∼2.5% at late-stage carcinoma), Foxp3^+^ T regulatory (Treg) cells (∼44% at early-stage and ∼68% at late-stage carcinoma), CD4^+^ T helper cells (∼20% at early-stage and ∼12% at late-stage carcinoma), cytotoxic CD8^+^ T cells (∼13.5% at early-stage and ∼6% at late-stage carcinoma), as well as CD19^+^ B cells (∼36% at early-stage and ∼44.5% at late-stage carcinoma) **(Fig. 1b)**.

To gain insights on the spatial distribution of the above populations in the tumor microenvironment, we conducted qualitative multiplex immunofluorescence analysis in MMTV-PyMT mice **(Figs. 1c-e)**, and indeed captured distinct subpopulations of CXCR4^+^PyMT^+^ tumor cells, CXCR4^+^CD4^+^ T cells, CXCR4^+^B220^+^ B Cells, and CXCR4^+^IBA1^+^ macrophages **(Figs. 1c-e)**. In consistency with our single cell data **(Fig. 1a)**, CXCR4 was restricted in either rare subsets of tumor cells **(Fig. 1e)** or the above-mentioned immune cells **(Fig 1c)**, but not in mesenchymal cells, e.g., PDGFRB^+^ pericytes/fibroblasts **(Fig.1d)**. Interestingly, all CXCR4^+^ cells were detected in relative proximity to EMCN^+^ blood vessels **(Figs. 1c-e)**, indicating spatial restriction and a perivascular trafficking pattern. Given that the main cell sources of CXCL12 according to our single cell data **(Fig. 1a)** and prior literature^15–20^, are endothelial cells and their associated perivascular mesenchyme, we hypothesized that the observed biased spatial distribution of the CXCR4^+^ cells was due to CXCL12 expression near the perivascular niches. To this end, we used multiplex immunofluorescence staining to capture intratumoral CXCL12 expression, and indeed found strict expression by EMCN^+^ endothelium and PDGFRB^+^ mesenchymal cells **(Fig. 1f)**.

When we investigated other cell populations, such as perivascular IBA1^+^ macrophages **(Fig. 1g)** and tumor cells **(Fig. 1h)**, we found either minimal, or background expression of CXCL12. Taken together, these data highlight that CXCL12 predominantly defines a perivascular niche within the breast cancer microenvironment, selectively attracting CXCR4^+^ populations, including distinct immune cell types and a subset of tumor cells, thereby establishing a spatially restricted pattern of CXCR4^+^ cell distribution near the CXCL12-expressing endothelial and mesenchymal cells.

To gain similar insights on human breast carcinomas, we mined publicly-available gene expression datasets, using The Cancer Genome Atlas (TCGA). In consistency with the mouse data **(Fig. 1)**, TCGA analysis confirmed that *Cxcr4* is significantly overexpressed in breast carcinomas, when compared to adjacent normal breast tissue **(Fig. 2a)**. Molecular subtype analysis using the widely accepted system for breast ductal adenocarcinoma classifying them into the Luminal, HER2^+^, and Triple Negative Breast Cancer (TNBC) subtypes, revealed increased *Cxcr4* expression in all three subtypes, with the latter exhibiting even higher *Cxcr4* expression compared to the Luminal and HER2^+^ subtypes **(Fig. 2a)**. Lastly, an investigation of *Cxcr4* expression in primary breast tumors compared to metastatic disease did not yield any difference **(Fig. 2a)**. These observations align with the prevailing view that the CXCR4^+^ breast tumor cell subset is preferentially selected in the primary tumor to undergo metastatic dissemination, facilitating its survival and proliferation at secondary sites^14^.

**Figure 2.**
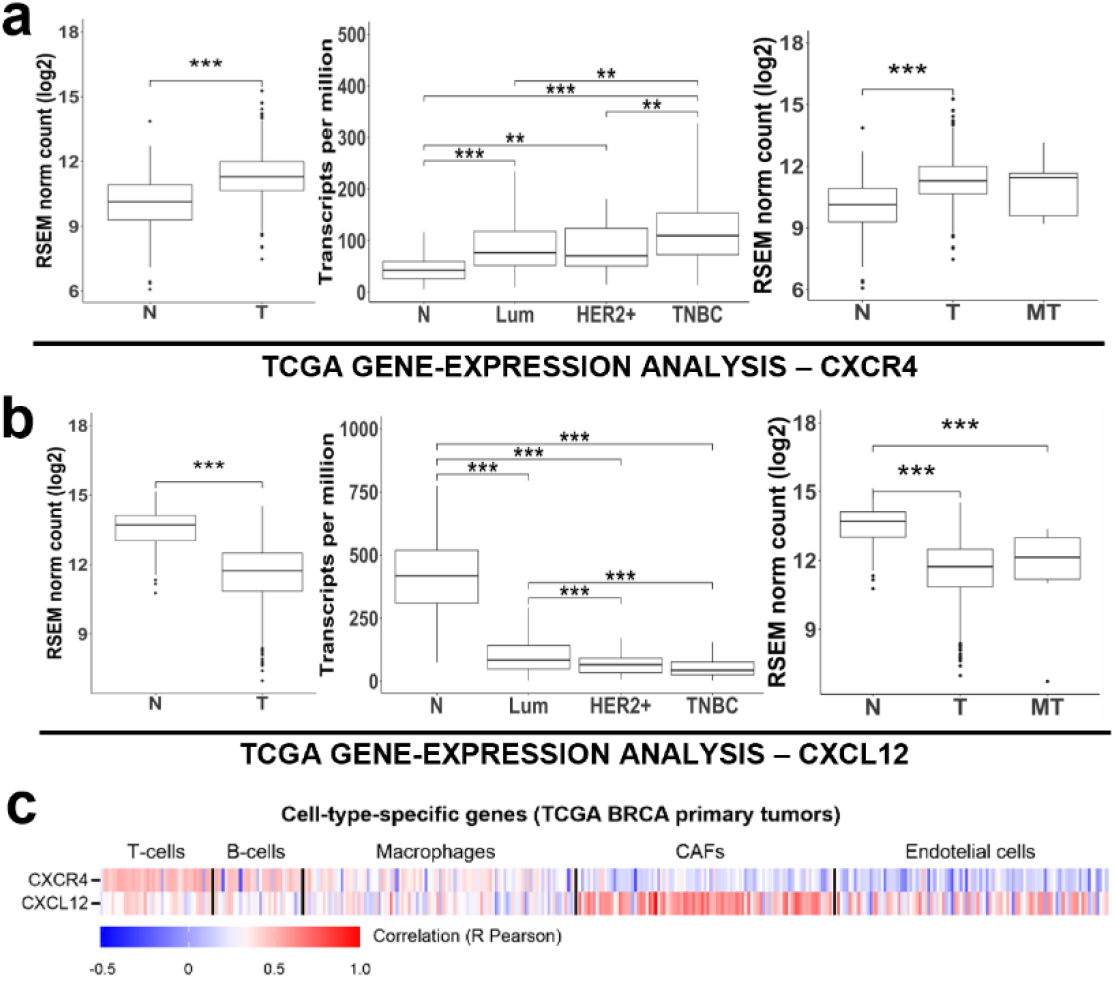
Expression of CXCL12 and CXCR4 in human breast cancer. (a-b) Publicly available transcriptomic TCGA data were used to compare normal (N), tumor (T) and metastatic tumor (MT) tissues of breast invasive carcinoma patients (Wilcoxon rank, \*\*\**p* < 0.001 for statistical significance). For gene expression analysis and comparison between the breast cancer subtypes, the UALCAN web platform with TCGA RNAseq data was used. Statistically significant gene expression differences between breast cancer subtypes [Luminal (Lum)], HER2^+^, Triple negative breast cancer (TNBC) and normal tissues (N) are indicated (\*\*\**p* < 0.001, ** P< 0.01, * P<0.05). **(c)** *In silico* analysis of tumor transcriptome from breast primary cancer samples (TCGA BRCA; n=1101). Hierarchical-Cluster heatmap reveals correlation of *Cxcr4* and *Cxcl12* gene expression with different cell types in all tumors. Cells are defined by sets of cell-type-specific genes (Supplementary Table 1).

In contrast, TCGA gene expression analysis for *Cxcl12* showed a significant decrease in *Cxcl12* levels in breast carcinomas, compared to the adjacent normal breast tissue **(Fig. 2b)**, with this decrease being consistent across all breast cancer molecular subtypes **(Fig. 2b)**. Moreover, no significant difference was found between primary and metastatic breast cancer, both showing lower *Cxcl12* levels, compared to normal breast tissue **(Fig. 2b)**. This might suggest that *Cxcl12* is downregulated in breast cancer; however, this interpretation overlooks the fact that *Cxcl12* is expressed by specific cell subsets within the tumor microenvironment, such as the mesenchymal and the endothelial cells, as demonstrated in our mouse model studies **(Fig. 1)**. As tumors grow, it is more likely that the increased ratio of tumor to stromal cells could dilute the overall *Cxcl12* expression. Thus, bulk tissue studies that combine epithelial and stromal components might falsely indicate downregulation of genes expressed by rarer cell subsets. To explore this possibility in more detail, we examined the co-expression of various chemokine gene networks with various cell type-specific gene signatures from TCGA breast tumors, categorized into “T cell-specific,” “B cell-specific,” “macrophage-specific,” “endothelium-specific,” and “mesen- chyme-specific” (“CAF”-specific) **(Fig. S4)**. Upon focusing on the CXCL12/CXCR4 pathway, we found that *Cxcl12* expression is primarily associated with the CAF- and endothelium-specific gene signatures **(Fig. 2c)**, thus suggesting that *Cxcl12* is contextually expressed in the perivascular niche, as in the case of murine tumors **(Fig. 1)**. Conversely, *Cxcr4* expression highly correlated with the macrophage- and lymphocyte-specific signatures, distinct from the endothelial and CAF associations **(Fig. 2c)**, again in consistency with the murine data **(Fig. 1)**.

Together, these data reveal a paracrine signaling interaction between the CXCL12^+^ perivascular niche and various CXCR4^+^ tumor and immune cell subsets in primary breast cancers.

### CXCL12/CXCR4 pathway suppression impairs metastatic dissemination in breast carcinoma

Only a few studies have previously investigated the expression and function of the CXCL12/CXCR4 pathway in the MMTV-PyMT mouse model^21,22^, and a mechanistic understanding of its role during the initial steps of the metastatic cascade is currently lacking. To address this issue, we utilized two models of suppression of the CXCL12/CXCR4 pathway. For the first method, we used the widely established CXCR4 antagonist, AMD3100, which competes against CXCL12 for ligand-binding to CXCR4, thus producing suboptimal CXCL12/CXCR4 responses within the tumor microenvironment^23–27^. In brief, MMTV-PyMT mice received 5 mg/Kg AMD3100 intraperitoneally daily for two weeks, before being subjected to various prometastatic readouts **(Fig. 3a)**. For the alternative method, we developed a CXCR4-deficient subline of the human metastatic MDA-MB-231 breast cancer cell line (231^shCxcr4^) along with the respective control vector-expressing subline (231^shCtrl^) **(Fig. S5a)**. We then used these sublines to generate orthotopic xenografts in immunodeficient SCID mice and assessed the same prometastatic readout after tumors reached ∼0.5 cm in size **(Fig. 2a’)**. As expected, the xenograft tumors resulted in cancer cell-specific elimination of CXCR4, while CXCR4 expression was disturbed for neither macrophages, nor endothelial cells **(Figs. S5b-c)**. Mice from both models were then subjected to prometastatic readouts **(Fig. S6a)**. In this regard, circulating tumor cells (CTCs) are the most direct prometastatic readout, because they are exclusively dependent on TMEM doorway function in breast cancer^28,29^. Indeed, we found a significant *(p<0.01; Mann-Whitney U-test)* ∼2-fold reduction of CTCs in AMD3100- treated as well as in 231^shCxcr4^ mice, when compared to their corresponding controls **(Figs. 3b-b’)**.

**Figure 3.**
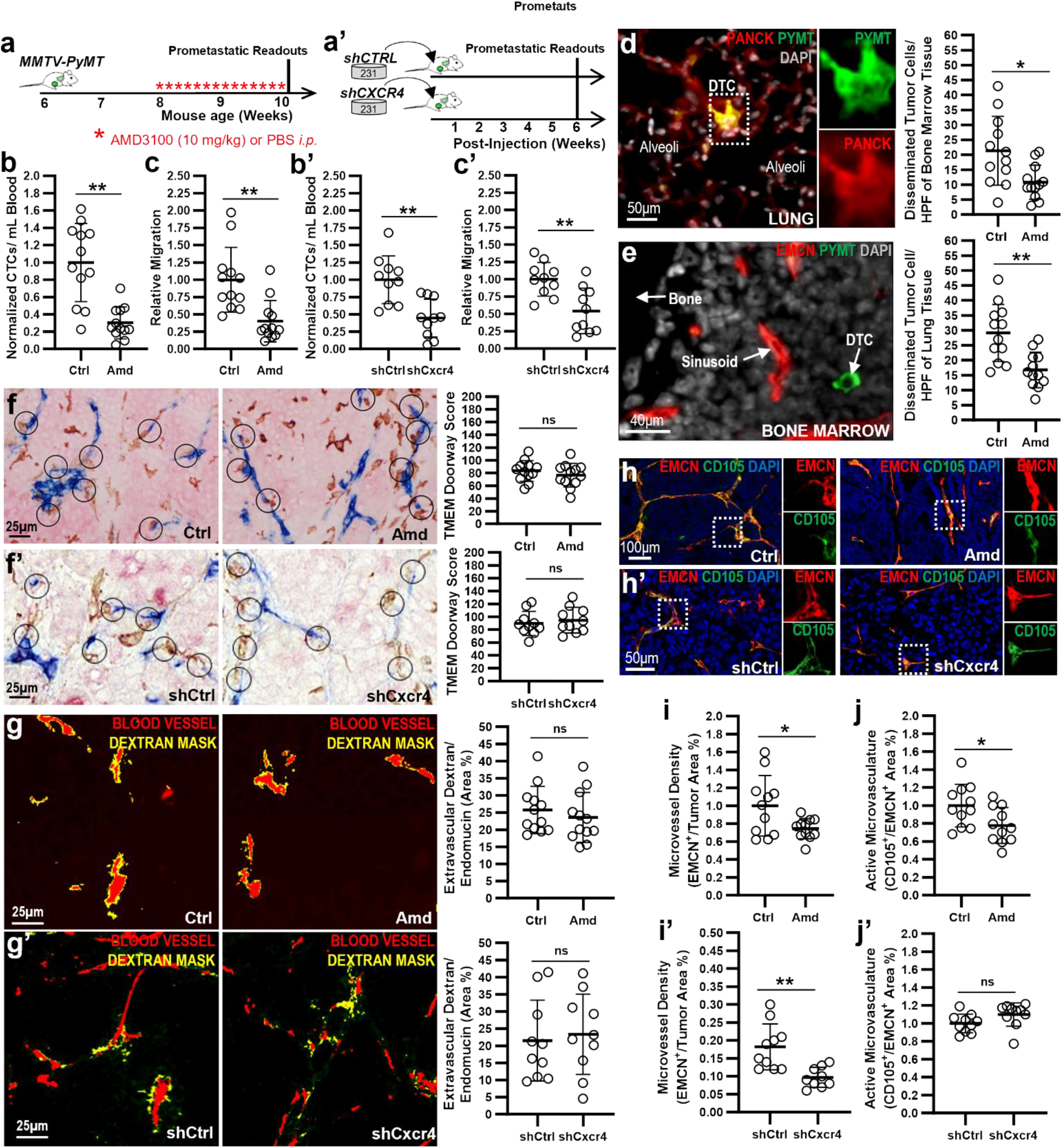
Suppression of the CXCL12/CXCR4 axis impairs cancer cell dissemination through disruption of promigratory signaling. (a-a’) Experimental pipeline for CXCL12/CXCR4 suppression in mouse models of breast carcinoma. In the first model, treatment with CXCR4 antagonist, AMD3100, began at ∼8 weeks in MMTV-PyMT mice and lasted for 2 weeks, after which mice were sacrificed and subjected to various prometastatic endpoints and assessments (a). In the second model, a CXCR4-null subline was generated from the parental MDA-MB-231 breast cancer cell line using shRNA, and was xenografted in SCID hosts, while control constructs (shCtrl) were used for comparison. The resulting tumors grew ∼1cm in diameter at ∼6-weeks post-injection, at which point, mice were sacrificed and sub- jected to various prometastatic endpoints and assessments (a’). **(b-b’)** Circulating tumor cells (CTCs) per milliliter of blood collected before sacrificing the mice in both models described in a-a’. Values nor- malized to the control group in each case, to account for inter-cohort variability. *Mann-Whitney U-test.* **(c-c’)** Relative migration of tumor cells passively entering CXCL12-coated needles, using the *in vivo* invasion assay in both models described in a-a’. *Mann-Whitney U-test.* **(d-e)** Disseminated tumor cells (DTCs) per surface area unit (mm^2^) of bone marrow tissue (d) or lung parenchyma (e), as assessed by the quantification of individual (magnified inserts) PyMT^+^ and/or PANCK^+^ tumor cells using multichannel immunofluorescence in the MMTV-PyMT mouse model. *Mann-Whitney U-test.* **(f-f’)** TMEM doorway identification by triple-stain immunohistochemistry (IHC) for each mouse model described in a-a’. Left panels: Representative images from each experimental condition. Right panels: TMEM doorway score, as assessed in 10 high-power fields (HPFs). *Mann-Whitney U-test.* **(g-g’)** TMEM doorway activity as- say, as assessed by the measurement of extravascular dextran leakage into the tumor parenchyma, following intravenous injection of Dextran-TMR before sacrificing the mice from both models described in (a-a’). Left panels: Representative blood vessel (endomucin) and extravascular dextran masks, as obtained by IF in mice, showing TMEM doorway-associated vascular permeability (yellow area). Right panels: Quantification of the surface covered by extravascular dextran, normalized to the blood vessel area in mice. *Mann-Whitney U-test.* **(h-j)** Evaluation of neoangiogenesis, using microvessel density and active microvasculature immunofluorescence pipelines in both models described in a-a’. Representa- tive images of multichannel immunofluorescence staining for endomucin (EMCN), CD105, and DAPI are illustrated for all experimental conditions (h). Microvessel density is reported as the area % cover- age of tumor parenchyma by EMCN^+^ blood vessels (i-i’). Active microvasculature is reported as the area % coverage of EMCN^+^ vasculature by CD105^+^ expression (j-j’). *Mann-Whitney U-test*.

We next sought to determine if the CXCL12/CXCR4 pathway is involved in blood-vessel directed migration of breast tumor cells^30^. Previously, we have developed an *in vivo* invasion assay that helps assess tumor cell invasion and migration through the passive collection of tumor cells migrating into chemoattractant-coated microneedles^31–33^. Using this setup **(Fig. S6b)**, we found that CXCL12-containing microneedles collected significantly *(p<0.001; Mann-Whitney U-test)* higher number of breast tumor cells, compared to PBS-containing microneedles **(Fig. S6b’)**, confirming that the *in vivo* invasion assay can be used to examine directional CXCL12-dependent migration. Interestingly, we found a significant *(p<0.01; Mann-Whitney U-test)* ∼2-fold reduction of CXCL12-mediated chemotactic migration of breast cancer cells into the microneedles in AMD3100-treated as well as in 231^shCxcr4^ mice, when compared to their corresponding controls **(Figs. 3c-c’)**.

As another useful prometastatic readout, we quantified single disseminated tumor cells (DTCs) in two frequent sites of breast cancer metastasis, lungs and bone marrow. In this regard, we performed multichannel immunofluorescence, by combining the pan-epithelial-specific antibody (pan-cytokeratin; PanCK), an antibody against the PyMT antigen capturing tumor cells, and a pan-endothelial-specific antibody (CD31) to demarcate vascular endothelia of the bone marrow and lungs **(Figs. S6c-d)**. In agreement, we found a significant *(p<0.01; Mann-Whitney U-test)* reduction of PanCK^+^PyMT^+^ DTCs in the bone marrow **(Fig. 3d)** and lungs **(Fig. 3e)** of MMTV-PyMT mice treated with AMD3100, when compared to their corresponding control.

We then examined whether the CXCL12/CXCR4 pathway has any direct effect on the assembly and/or function of the cancer cell intravasation, also known as Tumor Microenvironment of Metastasis (TMEM) doorways. First, to investigate TMEM doorway assembly, we performed a clinically-validated triple-stain TMEM immunohistochemistry assay^34–37^ that specifically demarcates the TMEM doorway cell triads, i.e., EMCN^+^ endothelial cell, IBA1^+^ macrophage and MENA^+^ tumor cell, in physical contact with one another **(Fig. S6e)**. This analysis revealed no significant *(p>0.05; Mann-Whitney u-test)* difference in TMEM doorway score for either of the models tested **(Figs. 3f-f’)**. Second, to investigate TMEM doorway function, we quantified TMEM doorway-specific vascular permeability, using a previously developed assay^38^ that measures TMEM doorway-mediated paracellular leakage of high-molecular weight dextran into the tumor parenchyma **(Fig. S6f)**. Using this assay, we found no significant *(p>0.05; Mann-Whitney U-test)* differences in extravascular dextran for either of the models tested **(Fig. 3g-g’)**. In support with the latter, we quantified disruption of TMEM doorway-associated endothelial junctions, by directly assessing the expression levels of the tight junction-specific protein zonula occludens-1 (ZO1) in tumor endothelia, along with extravascular dextran, using fixed-frozen tumor sections from MMTV-PyMT mice treated with either vehicle or AMD3100 **(Fig. S6g)**. In consistency, we found no significant *(p>0.05; Mann-Whitney U-test)* disruption of TMEM doorway-associated tight junctions between the two groups **(Figs. S6h-i)**. Along with the observations shown above **(Figs. 3a-e)**, these data additionally suggest that the CXCL12/CXCR4 pathway affects cancer cell intravasation by altering neither the assembly nor the activity of cancer cell intravasation (TMEM) doorways.

Given that CXCR4 is expressed by cancer-associated endothelium **(Fig. 1a)**, and this pathway is known to promote angiogenesis^39–41^, we examined whether CXCL12/CXCR4 suppression affected the tumor microvasculature in our models. Interestingly, Microvessel Density (MVD) was significantly reduced *(p>0.05; Mann-Whitney U-test)* following AMD3100 treatment **(Figs. 3h-i)** or genetic ablation of CXCR4 in tumor cells **(Figs. 3h’-i’)**, suggesting that the effects on angiogenesis might not be solely due to direct pathway involvement with endothelial cells. Supporting this, we assessed the “active” endothelium by measuring CD105 expression within the tumor microvasculature^42^ and documented significant reduction in CD105^+^EMCN^+^ endothelium with AMD3100 treatment *(p>0.05; Mann-Whitney U-test)* **(Figs. 3h&j)**. However, CXCR4 ablation in tumor cells did not impact blood vessel physiology **(Figs. 3h’&j’)**, again consistent with the premise that CXCL12/CXCR4 may elicit disparate mechanisms to affect endothelial homeostasis. Despite these findings aligning with the notion that CXCL12/CXCR4 affects neo-angiogenesis, we did not observe consistent changes in blood vessel morphology upon analyzing the vasculature network’s topography through skeleton analysis **(Figs. S7a-d & S7a’-d’)**.

In summary, CXCL12/CXCR4 appears to play a critical role in cancer cell dissemination by modulating specific biological programs within the metastatic cascade. Evidence suggests that this axis primarily influences the angiogenic potential of tumors and enhances the migratory capabilities of tumor cells. However, it does not seem to impact the biology of intravasation doorways, highlighting its selective contribution to certain stages of metastasis rather than a global effect on all metastatic processes.

### CXCL12 mediates translocation of prometastatic tumor cells toward TMEM doorways

Our data so far suggest that the CXCL12/CXCR4 pathway facilitates cancer cell dissemination from the primary tumor site without directly affecting the intravasation step of cancer metastasis, i.e., TMEM doorway formation and/or function. Because CXCL12 was consistently detected in endothelial cells, as well as in cells that are physically associated with the microvascular endothelium, such as in fibroblasts, and pericytes **(Figs. 1a and 1f)**, we hypothesized that CXCL12 could generate intratumoral gradients serving as chemotactic forces for the translocation of CXCR4^+^ tumor and other cells to the perivascular niche, and more specifically, the TMEM doorways. We thus examined if CXCL12 expression is spatially correlated with intratumoral blood vessels and/or TMEM doorways. To address this, we developed a new image analysis method, in which the expression level of any protein, in this case CXCL12, can be progressively quantified in concentric annuli, drawn in consistent interval areas from designated regions of interest (ROIs). To investigate the spatial correlation of CXCL12 with TMEM doorways, we first aligned the tumor section stained via TMEM doorway IHC with a corresponding sequential section stained with endomucin (EMCN; blood vessel identification) and CXCL12, using immunofluorescence (IF) staining **(Figs. 4a-a’)**, and visually confirmed that CXCL12 was highly expressed in the immediate surroundings of TMEM doorways, thus revealing an “asymmetric” pattern of expression around the blood vessels **(Figs. 4a-a’)**. TMEM doorway ROIs were then demarcated on the IHC section through vector-based analysis, and aligned with the sequential IF sections **(Figs. 4b-b’)**. In the sequential IF section, concentric annuli were then drawn as new independent ROIs spanning away from the TMEM doorways **(Fig. 4c)**. The EMCN channel served as an exclusion mask in this analysis, to ensure that CXCL12 expression would only be measured within the tumor parenchyma, and not intravascularly **(Fig. 4d)**. The generation of these annulus-shaped ROIs allowed us to measure cumulative CXCL12 expression (green mask) along progressively increasing distances from TMEM doorways **(Fig. 4e)**. In addition, to measure CXCL12 expression gradients generally spanning from the perivascular niche (PN), and not necessarily from the TMEM doorways, we simply generated concentric annuli around the entire blood vessel using the EMCN stain as the designated ROI **(Fig. 4c’-e’)**. Quantification of the CXCL12 signal within individual annuli around TMEM- or PN-ROIs revealed a stronger *(p<0.05; two-way ANOVA)* intratumoral CXCL12 gradient around TMEM doorways, when compared to PNs **(Fig. 4f)**. Interestingly, CXCL12 gradients spanned along a ∼45µm distance when assessed from their respective TMEM doorways, but plateaued in shorter distances, when assessed in entire blood vessels **(Fig. 4f)**.

**Figure 4.**
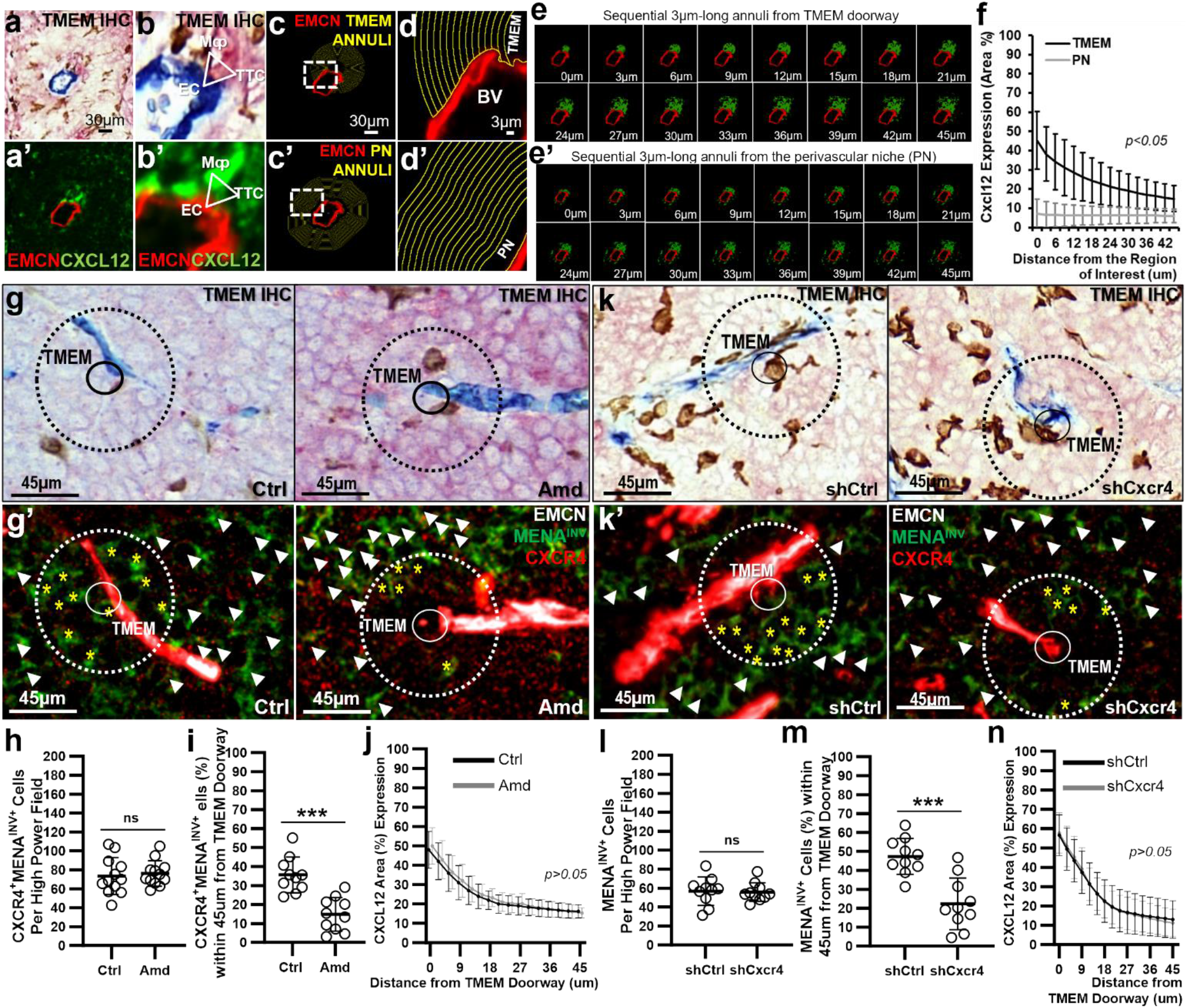
CXCL12 forms an intratumoral gradient around perivascular cancer cell intravasation doorways, to chemotactically attract CXCR4^+^ tumor cells. (a-a) Co-registration of sequential slides from PyMT animals stained for either TMEM doorway triple-immunohistochemistry (a), or multichannel immunofluorescence for endomucin (EMCN) and CXCL12 (a’). The EMCN signal signifies the same vascular profile in the middle of each image. **(b-b)** Magnified inserts focused on TMEM doorway region from the corresponding images on (a-a’). The triangular vector pinpoints to the triad of cells composing TMEM doorways: EC, endothelial cell; TTC, TMEM Tumor Cell; Mφ, macrophage. **(c-c’)** Development of concentric annuli circumnavigating around either a TMEM doorway region-of-interest (TMEM annuli; c), or a perivascular space region-of-interest (PN Annuli, c’). **(d-d’)** Magnified inserts of the white-dashed boxes shown in (c-c’), revealing the nature of concentric annuli around each corresponding region of interest, with the luminant component of the blood vessel, being excluded from the annuli. **(e-e’)** Serial images revealing CXCL12 signal mask in each sequential annulus, from the designated regions-of-interest shown in (c-c’). **(f)** Quantification of the area % covered by CXCL12 signal within each individual annulus starting from those closest to the designated region-of-interest, i.e., TMEM doorway (c, d, e), or the PN (c’, d’, e’), to those at 45um away from the region-of-interest. Graph shows the average CXCL12 intensity in each annulus from 30 vascular profiles **(g-g’)** Co-registration of sequential slides stained for either TMEM doorway triple-immunohistochemistry (g), or multichannel immunofluorescence for endomucin (EMCN), MenaINV, and CXCR4, in MMTV-PyMT mice treated with either vehicle (1^st^ column) or Amd3100 (2^nd^ column). Small circles represent vectors pinpointing TMEM doorways in both sections. Large-dotted circles represent the TMEM doorway “area of influence”, which spans 45um away from the TMEM doorway, as demonstrated in (f). Yellow asterisks depict CXCR4^+^MENA^INV+^ tumor cells within the TMEM doorway “area of influence”. White arrowheads depict CXCR4^+^MENA^INV+^ tumor cells away from the TMEM doorway “area of influence”. **(h)** Quantification of the total number of CXCR4^+^MENA^INV+^ tumor cells per high-power field in MMTV-PyMT mice treated with either vehicle, or AMD3100, irrespective of their positioning relative to the TMEM doorway area of influence. *Mann-Whitney U-test.* (i) Quantification of CXCR4^+^MENA^INV+^ tumor cells in MMTV-PyMT mice treated with either vehicle, or AMD3100, exclusively within the TMEM doorway area of influence. *Mann-Whitney U-test. (j) Assessment of CXCL12 gradient around TMEM doorways, as described in (f), in MMTV-PyMT mice treated with either vehicle, or AMD3100.* (k-n) Identical analyses to those described in (g-j), using the 231^shCtrl^/231^shCxcr4^ mouse xenografts.

Given the high expression of CXCL12 in endothelial cells and various perivascular cells **(Figs. 1a and 1f-h)**, it is expected that intratumoral CXCL12 gradients are concentrated within the perivascular niche (PN), particularly in TMEM doorways. These doorways represent “specialized” sub-compartments for intravasation within the PN. Because TMEM doorways lack peri-endothelial pericytes^29^, we hypothesized that the CXCL12 gradient in these regions primarily originates from the endothelium, associated with TMEM doorways and the peripheral blood. TMEM doorways facilitate metastatic dissemination by providing openings to the endothelial vasculature^29^. To investigate whether TMEM doorway function contributes to CXCL12 influx in the breast cancer microenvironment, we utilized the Macrophage Fas-Induced Apoptosis (MaFIA) mouse model, which allows the inducible and reversible apoptosis of macrophages and dendritic cells upon B/B homodimer administration^43^. Tumors from syngeneic 12-week-old MMTV-PyMT hosts were transplanted into MaFIA mice, which were then treated with the B/B homodimer or vehicle control once tumors reached approximately 0.5 cm in size **(Fig. S8a)**. Macrophage depletion was observed to be about 75% in the B/B homodimer-treated mice compared to the vehicle-treated controls **(Fig. S8b-c)**. Previous research indicated that macrophage depletion in MaFIA mice disrupts TMEM doorway function^44^, suggesting that B/B administration should reduce intratumoral CXCL12 gradients if they indeed originated from TMEM doorway-mediated vascular permeability. Our analysis of CXCL12 expression after eliminating the intraluminal CXCL12 signal using a blood vessel mask, revealed significantly *(p>0.05; Mann-Whitney U-test)* lower CXCL12 levels in MaFIA-PyMT mice with macrophage depletion (B/B homodimer group) compared to controls **(Figs. S8d-e)**. Additionally, there was a significant *(p<0.01; Mann-Whitney U-test)* decrease in the number of vascular “hotspots” associated with TMEM-doorway CXCL12 gradients **(Figs. S8d&f)**. Together, these findings suggest that CXCL12 is likely derived from the blood plasma, and that macrophage depletion impairs TMEM doorway-associated vascular openings, thereby confining CXCL12 to the blood plasma and preventing its leakage into the tumor parenchyma.

Chemotactic signals originating from the tumor microvasculature, such as the hepatocyte growth factor (HGF), have been shown to attract promigratory MENA^INV+^ tumor cells from as far as 500 µm away from the blood vessel source. However, HGF is not specific to TMEM doorways^30^ as CXCL12 is, based on our findings. Thus, we hypothesized that while promigratory MENA^INV+^ tumor cells respond to HGF gradients over long distances within the tumor microenvironment to be generally attracted to blood vessels^30^, they are potentially redirected specifically towards a TMEM doorway if they move into a CXCL12 gradient within ∼45 µm distance from the doorway. In support with this hypothesis, we first confirmed that the promigratory MENA^INV+^ tumor cells co-express CXCR4 in PyMT mice **(Figs. S9a-c)**. The pharmacological inhibition of the CXCL12/CXCR4 pathway using AMD3100 did not significantly *(p>0.05; Mann-Whitney U-test)* alter the total number of CXCR4^+^MENA^INV+^ tumor cells **(Figs. 4g-g’&h)**. However, AMD3100 significantly *(p<0.01; Mann-Whitney U-test)* reduced the CXCR4^+^MENA^INV+^ breast cancer cells within the 45 µm-radial ROIs spanning from TMEM doorways **(Figs.4g-g’&i)**. Importantly, TMEM-doorway generated CXCL12 gradients were not affected by AMD3100 treatment **(Fig. 4j)**, thus indicating that the altered localization of CXCR4^+^MENA^INV+^ tumor cells is purely due to the antagonistic functions of AMD3100 on CXCR4^+^ tumor cells, and not due to modifications in the CXCL12 gradient itself. Finally, we were able to confirm similar observations in the xenogeneic 231^shCxcr4^/231^shCtrl^ model, although in this case, the promigratory tumor cells were assessed exclusively on the basis of MENA^INV^ expression, because CXCR4 has been genetically knocked down in tumor cells **(Figs. 4k-n)**.

Collectively, these data suggest that TMEM doorways attract via chemotaxis the dissemination- competent tumor cells in a CXCL12/CXCR4-dependent manner.

### The intravasation doorways orchestrate an immunosuppressive milieu in breast carcinomas

We previously proposed that the regulation of immunosuppressive tumor microenvironments may be context-dependent and suggested that TMEM doorways could represent immunosuppressive niches^45^. Given that CXCL12 is a well-established immunosuppressive chemokine^46–49^, with its expression being restricted around TMEM doorways **(Figs. 4a-f)**, and the fact that multiple immune cell subsets, in addition to tumor cells, seem to express CXCR4 **(Fig. 1)**, we hypothesized that TMEM doorways might orchestrate localized immunosuppression through the CXCL12/CXCR4 axis. To explore whether TMEM doorways could indeed represent immunosuppressive niches, we employed a digital pathology approach using Multiple Immunohistochemistry Consecutive Staining on a Single Slide (MICSSS)^50^ targeting a panel of 10 lineage and functional immunomarkers in 6 breast cancer patients with invasive breast carcinomas. As this panel has been optimized for clinical use in human patients, the traditional markers were not suitable for detecting TMEM doorways. Specifically, MENA, which is typically used to detect tumor cells in the TMEM doorway IHC assay, was replaced by pancytokeratin (PANCK), while the endothelial (CD31) and macrophage (CD68) markers remained unchanged. To this end, we aligned individual images in the same fields-of-view (FOVs), and cell triads composed of a PANCK^+^ tumor cell, a CD31^+^ endothelial cell and a CD68^+^ macrophage, using the CD31-CD68-PANCK composite image were reconstructed in QuPath, herewith called “intravasation portals”. Matched peri- vascular niches lacking CD68^+^ macrophages, but retaining both the PANCK^+^ tumor cells and the CD31^+^ endothelial cells, were selected as “control” regions-of-interest for comparison. The remaining markers from the MICSS panel were used to assess the relative abundance of various immune cell subsets and their functionality within the surrounding ROI from intravasation portals or control regions, in automated fashion. We found an increased relative proportion of CD8^+^ T cells within perivascular niches containing intravasation portals, when compared to those lacking intravasation portals **(Figs. 5a-b)**. However, a significant proportion of these CD8^+^ T cells were identified as GZMB^+^ and PD1^+^ **(Figs. 5a-b)**, indicating that perivascular niches with intravasation portals are more likely to harbor CD8^+^ T cells lacking Granzyme-B expression and overexpressing immune checkpoint receptors, like PD1, both indicative of reduced effector functions. This finding suggests that while the trafficking into and out of intravasation portals is not notably restricted, T cells that enter these regions often present with diminished activity.

**Figure 5.**
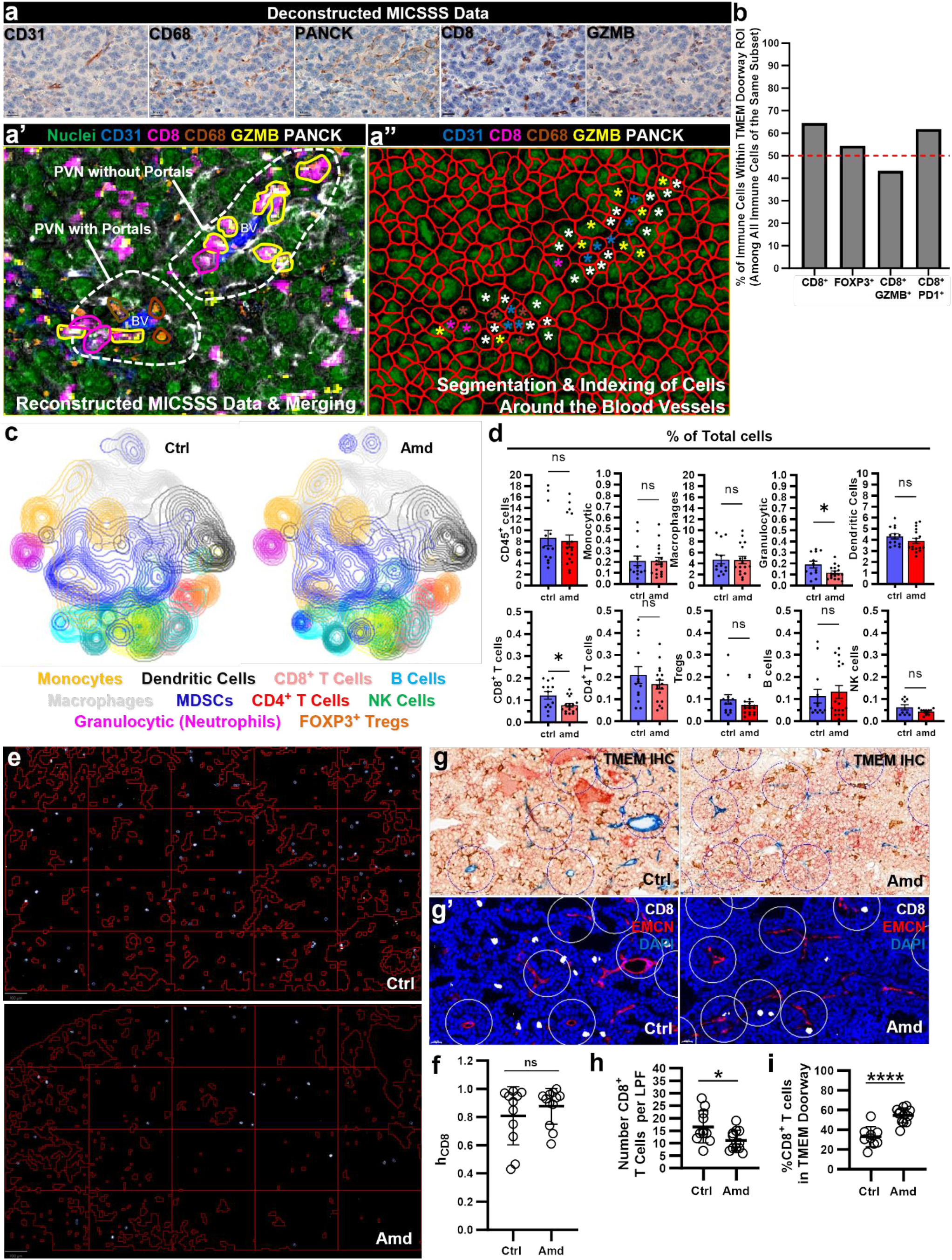
TMEM doorways form an immunosuppressive niche in the tumor microenvironment. (a-a’’) Multiple Immunohistochemistry Consecutive Staining on a Single Slide (MICSSS) analysis, to quantify the presence of immune cell subsets and their functional states. The deconstructed MICSSS images of 5 individual markers, CD31, CD68, PANCK, CD8, GZMB, are shown in (a), after co-registering them into the same field-of-view. These channels were reconstructed into a single image, shown in (a’), by pseudo-coloring to distinguish between the individual signals. The hematoxylin channel labeling nuclei was pseudo-colored in green, to facilitate cell segmentation, as described in the main text and materials and methods. Using classifiers, we identified cell triads in close/triangular physical proximity within the field-of-view, corresponding to CD31^+^ endothelial cells, CD68^+^ macrophages, and PANCK^+^ tumor cells, representing putative intravasation portals. Note that because the MICSSS panel was pre- fixed and not customized, MENA, which is traditionally used in TMEM-IHC to identify TMEM doorway-associated tumor cells was not used here, and was replaced by PANCK. Perivascular spaces lacking CD68^+^ macrophages were used as control regions-of-interest designated as lacking the intravasation portals. Immune cell subsets were then quantified within and compared within these different niches. Yellow-encircled cells are CD8^+^ T cells with high GZMB expression, while magenta-encircled cells are CD8^+^ T cells lacking GZMB expression. Cell identities were tracked within this niches using classifier tools, and assigned a unique identity, pointed for simplicity in (a’’) with a unique color asterisk. **(b)** Relative proportions of immune cell subsets within intravasation portal niches. Interestingly, 60% of CD8^+^ T cells are preferentially located within the intravasation portal niche, although only 40% of CD8^+^ T cells within these niches are GZMB^+^, indicating that despite their biased chemo-attraction toward intravasation portals, CD8^+^ T cells present with lower effector functions. In conjunction with this, the relative proportion of CD8^+^ T cells expressing the inhibitory checkpoint receptor PD1 is also enriched, at 60% of total CD8^+^PD1^+^ T cells. Although this study was focused on CD8^+^ T cells, it should be reminded that MICSSS panel consisted of multiple markers, enabling the quantification of other immune cell subsets, for example FOXP3^+^ T regulatory cells, also shown here, which did not reveal any difference in distribution between the two niches. **(c)** T-SNE overlays of subsets of immune cells as defined by color codes in MMTV-PyMT mice treated with either vehicle or Amd3100, using multiparametric flow cytometry. **(d)** Quantification of the individual immune cell subsets in MMTV-PyMT mice treated with either vehicle or Amd3100, using flow cytometry. *Mann-Whitney U-test.* **(e-f)** Assessment of homogeneity (h) of CD8^+^ T cell distribution in the tumor microenvironment. Representative images from MMT’V-PyMT mice treated with either vehicle (Ctrl), or Amd3100 (Amd) are shown (e). Bright, cyan-labeled cells are CD8+ T cells. Vertical and horizontal straight red lines define individual sectors in the tumor microenvironment. Regions enclosed in the remaining continuous red lines are excluded ROIs corresponding to necrotic areas, blood vessel lumens or non-tumor tissues. Homogeneity is assessed by comparing the deviation of CD8+ T cell densities among the sectors. Quantification of the homogeneity of CD8^+^ T cell distribution (h_CD8_) between vehicle-treated and Amd3100-treated PyMT mice is shown in (f). Mann-Whitney U-test. **(g-i)** Co-registration of sequential slides stained for either TMEM doorway triple-immunohistochemistry (g), or multichannel immunofluorescence for endomucin (EMCN), and CD8, in MMTV-PyMT mice treated with either vehicle (1^st^ column) or Amd3100 (2^nd^ column) (g’). White circles represent the TMEM doorway “area of influence”, which spans 45um away from the TMEM doorway, as demonstrated in Figure 4. The number of CD8^+^ T cells was quantified either in the entire low power field, revealing the significant decrease of CD8^+^ T cell density in the tumor microenvironment overall, following treatment with Amd3100 (h), or exclusively within the TMEM doorway area of influence (white circles), revealing a biased increase in the distribution of CD8^+^ T cells near the TMEM doorways, following treatment with Amd3100 (i). *Mann-Whitney U-test*.

While more comprehensive analyses of immune cell subsets using our MICSSS platform was beyond the scope of this article, these observations collectively support the notion that TMEM doorways may function as immunosuppressive niches, limiting CD8^+^ T cell function, in human breast cancer patients.

We subsequently sought to determine if the CXCL12/CXCR4 pathway regulates the composition of immune cells in the breast tumor microenvironment. First, we examined the general influx of multiple immune cell populations in MMTV-PyMT mice treated with AMD3100 or vehicle-control. We gated cells based on forward and side scatter, excluded cell doublets, and focused on the CD45^+^ hematopoietic-derived immune cell population **(Fig. S3)**. Next, we used t-Distributed Stochastic Neighbor Embedding (t-SNE) for unsupervised clustering and dimensionality reduction, and the identification of distinct or similar clusters of cells and their relative proportions. The t-SNE analysis and population quantification across individual mice revealed no significant changes in most immune cell subsets examined, but there was a minor disruption with the numbers of granulocytic (neutrophil) cells, and CD8^+^ T cells in AMD3100-treated mice **(Figs. 5c-d)**. However, upon inspecting the expression of a wide variety of functional markers included in this high-dimensional flow cytometry panel, we failed to uncover any shifts on the phenotypes of CD8^+^ T cells **(Fig. S10a)**, CD4^+^ T helper cells **(Fig. S10b)**, FOXP3^+^ Tregs **(Fig. S10c)**, NK-T cells **(Fig. S10d)**, B cells **(Fig. S10e)**, and F4/80^+^ macrophages **(Fig. S10f-g)**. Neutrophils and other granulocytic myeloid cells are generally known to express CXCR4 and to constitute an immunosuppressive cell population within the tumor microenvironment^41,45,51^. On the contrary, the trafficking of the cytotoxic CD8^+^ T cells into the tumor microenvironment relies on CXCL9/10/11 and CCL5^52–57^, instead of CXCL12, suggesting that the mild trafficking defect could be indirect, and even possibly attributable to the observed granulocyte trafficking impairment. Although other immune populations, such as F4/80^+^ macrophages and B cells do express CXCR4, their chemotactic activity is complex and may be predominantly dependent on other chemokine axes, such as CCL2/CCR2^58–61^ and CXCL13/CXCR5^62–65^, respectively, thus explaining why AMD3100 may not have directly impaired their influx into the tumors. Together, these findings indicate that while the CXCL12/CXCR4 pathway minimally affects the composition of most immune cell subsets in the breast tumor microenvironment, it appears to mildly influence the trafficking of granulocytic cells and potentially of cytotoxic CD8^+^ T cells.

Because there were no significant shifts in overall immune cell composition after CXCL12/CXCR4 suppression **(Figs. 5c-d)**, and given that CXCL12 expression is limited around the TMEM doorway **(Figs.4a-f)**, we theorized that the CXCL12/CXCR4 pathway regulates immune cell distribution within the tumor microenvironment after entering the primary tumor microenvironment. To address this, we employed two independent, supervised methods to assess CD8^+^ T cell topology in MMTV-PyMT mice treated with either AMD3100 or vehicle. In the first method, we conducted a CD8^+^ T cell homogeneity analysis using low-power field to examine CD8^+^ T cell distribution on a global scale **(Figs. S11a-c)**.

This approach will reveal whether only specific tumor compartments are trafficked by CD8^+^ T cells, or whether the CD8^+^ T cell distribution is more uniform, thus helping determine whether the tumors exhibit characteristics of immune-cold or immune-hot composition at a macro-scale. In the second method, we carried out a sub-analysis of CD8^+^ T cell distribution on high-power fields, via annulus morphometry around TMEM doorways. Given the restricted expression of CXCL12 around the TMEM doorways, this method will examine if the CXCL12/CXCR4 pathway finetunes CD8^+^ T cell distribution toward or away from these perivascular portals, thus providing insight into the localized effects of CXCL12 on immune cell positioning in relation to the vasculature. As expected, there was no significant difference *(p>0.05; Mann-Whitney U-test)* in the homogeneity of CD8^+^ T cell distribution when assessed at the macroscale across the entire tumor (method 1) **(Figs. 5e-f)**, indicating that CD8^+^ T cells are capable of infiltrating all tumor regions uniformly. However, when we analyzed CD8^+^ T cell distribution in relation to the TMEM doorways using annulus morphometry (method 2), we observed *increased (p<0.001; Mann-Whitney U-test)* trafficking of CD8^+^ T cells within the 45-µm radius around the TMEM doorways. Notably, while this localized trafficking was evident, the overall density of CD8^+^ T cells was slightly reduced *(p<0.05; Mann-Whitney U-test)* following AMD3100 treatment **(Figs. 5g-i)**, a finding that was further supported by the flow cytometry analysis **(Fig. 5d)**.

In summary, the data presented in this section collectively suggest that the CXCL12/CXCR4 axis does not only influence the overall distribution of CD8^+^ T cells, but also enhances their localization to specific tumor microenvironmental features, such as TMEM doorways, despite an overall decrease in their numbers.

## DISCUSSION

It has been long established that CXCL12/CXCR4 is implicated in breast cancer metastasis^9,12,20,66^, but detailed mechanistic understanding of the process during the initial steps of the metastatic cascade has been lacking. Here, we demonstrated that CXCL12 expression is not homogeneously distributed within the primary breast tumor microenvironment, but instead, it is expressed as a gradient around cancer cell intravasation doorways, also known as TMEM doorways, thus serving as a strong chemotactic factor for the translocation of intravasation-competent (CXCR4^+^MENA^INV+^) tumor cells to the peri- vascular space, whereby they can find entry into the circulation via TMEM doorways. Indeed, both the genetic ablation and the pharmacological suppression of the CXCL12/CXCR4 pathway, resulted in significantly impaired translocation of the CXCR4^+^MENA^INV+^ tumor cells to TMEM doorways, leading to impairment of cancer cell dissemination in the peripheral circulation. Although it has been previously postulated that CXCL12 is upregulated in perivascular regions within tumors, and is responsible for chemotaxis of tumor-associated macrophages (TAMs)^19^, here, we have shown for the first time that perivascular CXCL12 expression is even more strictly contextualized at the perivascular niche, and more specifically, it is spatially linked to the TMEM doorways.

Our study has revealed the mechanism by which chemotactic ligands control migration of MENA^INV+^ tumor cells to TMEM doorways. A requirement for directional cancer cell migration is asymmetric actin polymerization in response to a chemotactic gradient, resulting in cancer cell locomotion towards the chemoattractant source^67,68^. In particular, HGF acts as long-distance chemoattractant for cancer cells towards blood vessels^30^, while co-migration of macrophages with tumor cells to TMEM doorways resulting from the EGF-CSF1 paracrine signaling loop, operates at distances shorter than 500µm^32,69^. Cancer cells capable of responding to these chemotactic signals typically express MENA^INV^, which enhances sensitivity of receptor tyrosine kinases to their ligands^70–72^. As such, the combination of EGF and HGF chemotactic signals in the tumor microenvironment maintains, on one hand, the tumor cell-macrophage pairing [which in turn promotes the induction of MENA^INV^ in the tumor cells^73^], while on the other hand, facilitates their coordinated migration towards the perivascular niche. The novel insight from our study is that when tumor cells reach approximately 45µm away from the blood vessels, CXCL12 signaling biases the migration of MENA^INV+^CXCR4^+^ tumor cells toward TMEM doorways, which facilitates transendothelial migration. It is likely that migratory cells that fail to detect this “narrow” peri-TMEM doorway CXCL12 gradient, will also translocate to the perivascular niche under the control of the HGF gradient, but they will never intravasate as they will be located away from functional TMEM doorways. Thus, our observations provide an attractive model of the molecular interplay between the HGF and CXCL12 chemokines in the tumor microenvironment.

Although our study has clearly demonstrated that invasive/migratory CXCR4^+^/MENA^INV+^ tumor cells could translocate to TMEM doorways under the controlled regulation of the newly described TMEM-derived CXCL12 expression gradient, and thus facilitating cancer cell intravasation and subsequent metastatic dissemination, we should bear in mind that the CXCL12/CXCR4 pathway may be negatively or positively regulated by a plethora of other microenvironmental factors. Although initially described as an orphan receptor^74^, the atypical chemokine receptor-3 (ACKR3), later embraced into the chemokine receptor family as CXCR7, has been recognized for a long time as a second receptor for CXCL12^75^. Prior reports have demonstrated that CXCR7/ACKR3 is frequently upregulated in both tumor cells and tu- mor-associated endothelium in breast cancer^76,77^. In particular, CXCR7/ACKR3 prevents breast cancer metastasis via reducing cancer cell intravasation and via promoting anti-metastatic endothelial properties, thus opposing the functions of CXCL12/CXCR4^16,66,78^. In line with the above, CXCR7/ACKR3 may enact a major regulatory function on the CXCL12/CXCR4 signaling axis in cancer cell dissemination pathway, potentially operating as a scavenger receptor that binds to, and internalizes CXCL12, thus directly modifying ligand bioavailability in the tumor microenvironment^79–82^. Future research should focus on further explorations of CXCR7/ACKR3 in regulating CXCL12/CXCR4-dependent translocation of prometastatic tumor cells to TMEM doorways.

Besides cancer cell migration and metastasis, CXCL12 may act as an orchestrator of multiple other cancer hallmarks, such as angiogenesis^83^, induction and maintenance of the cancer stem cell niche^84^, as well as induction of an immunosuppressive tumor microenvironment^15,17,85^. Especially for the latter, it seems that the role of CXCL12 is intricate and not as clear. For instance, an early study revealed that cytotoxic T cells are chemotactically attracted by low concentrations of CXCL12, but repelled by high concentrations of CXCL12^86^. It was specifically shown that although both behaviors are regulated via CXCL12/CXCR4-dependent signaling, there are key differences in the propagation of the intracellular signaling relay, i.e., the former was PI3K-dependent, while the latter behavior was cAMP-dependent^86^. In oral squamous cell carcinoma, increased CXCL12 expression levels positively correlate with FOXP3^+^ T regulatory cell infiltration into the tumor, but inversely correlate with cytotoxic CD8^+^ T cell infiltration, again confirming the phenotypic heterogeneity^49^. Besides the direct effects of CXCL12/CXCR4 on various T cell subsets, myeloid cell populations, such as tumor-associated macrophages (TAMs) and mye- loid-derived suppressor cells (MDSCs), are also known to highly express CXCR4, and they can home into tumors to confer immunosuppressive niches^45,87^. These intricacies indicate that both intrinsic factors and microenvironmental cues come at play to determine the overall outcomes of CXCL12/CXCR4 in immune cell chemotaxis. The current work is thus consistent with the paradigm that CXCL12 may function as a general chemoattractant by bringing CXCR4^+^ immune cells into the tumor and specifically around the TMEM doorways. However, CD8^+^ T cells attracted in these niches were immunosuppressed in human patients (i.e., PD1^+^GZMB^-^) indicating that T cells with reduced effector functions may develop either directly from CXCL12/CXCR4-mediated signaling, or due to the unique composition of the tumor microenvironment in TMEM doorways, such as presence of perivascular/proangiogenic macrophages. Although multiple immune cell populations were found to express CXCR4, our studies also indicate that CXCR4^+^ cells are often minority subsets. For example, only ∼10% of total CD8^+^ T cells were found to be CXCR4^high^, based on flow cytometry. This finding complicates the interpretation of the trafficking patterns observed after administration of the CXCR4 antagonist, AMD3100, which reveals a significant reduction of CD8^+^ T cells in the tumors. There are two possible explanations for this phenotype. First, chemotactic pathways typically present with very low signaling thresholds^67,88^, and as such, it is possible that even CXCR4^low^ CD8^+^ T cell populations could be responsive to intratumoral CXCL12 gradients. Second, due to the high heterogeneity of CXCR4^+^ immune cell subsets in the tumor microenvironment, it is also likely that the reduced influx of CD8^+^ T cells into the tumor parenchyma could be an indirect result of disturbing other immune cell populations such as myeloid cells, which in turn fail to attract CD8^+^ T cells through T cell-specific chemokines, such as CXCL9/10. As such, chemotactic cascades and context-dependent synergistic networks may obfuscate T cell trafficking into the tumors.

Of particular interest is the CXCL12/CXCR4 pathway, often viewed as an archetypal homing chemokine mechanism^41^, yet not considered a primary chemotactic pathway for most immune cell subsets. For example, T cells primarily use the CXCL9/10/11-CXCR3 axis^89–91^, while B cells rely mostly on the CXCL13-CXCR5^63^ axis, and macrophages on the CCL2-CCR2 axis^61^. Thus, CXCL12/CXCR4 may serve as a "fine-tuning" mechanism that influences the spatial positioning of immune cells within the diverse intratumoral niches once they enter the tumor^41^. Our large-scale homogeneity analyses did not reveal significant changes in CD8^+^ T cell distribution in the tumor microenvironment following AMD3100 administration. However, when we focused on cellular resolution, we observed misplacement of CD8^+^ T cells relative to the perivascular space and TMEM doorways. These findings suggest that the CXCL12/CXCR4 pathway acts as an auxiliary mechanism, providing specific guidance cues for tumor and immune cell populations in relation to cancer vasculature and intravasation sites, without impacting their initial presence in the tumor. This may also partially explain the suboptimal outcomes of therapies targeting the CXCL12/CXCR4 pathway in clinical trials.

Moving forward, our work underscores the importance of viewing chemokine pathways, in particular CXCL12/CXCR4, in a more holistic manner, specifically as synergistic networks. These networks may significantly influence the migratory behavior of tumor and immune cells in relation to the perivascular space. By gaining a comprehensive understanding of the functional prerequisites that drive preferential attachment of prometastatic tumor cells and immunosuppressive immune cells to the perivascular niche, we can achieve better control over metastatic dissemination, and potentially prevent it altogether through pharmacological interventions.

## AUTHOR CONTRIBUTIONS

D.P.A. and G.S.K. conceived the project; D.P.A, P.S.F., V.P.L., J.S.C., R.R., H.G.H., A.L.M, E.R.T, L.M.L., G.L., and G.S.K. designed experiments. D.P.A., N.C., S.G., D.G.A., S.V., L.R.S., E.G., J.K., A.G., A.K., J.B., N.R., C.L.D., A.S.H., S.S., D.E., A.Z., X.C., R.J.E., M.H.O., G.I., S.G., A.B., H.G., R.A., V.D., P.S.F., V.P.L., A.Q., D.M.R., and R.G, performed all the experiments and conducted bioinformatic and data analyses. D.P.A and G.S.K wrote the manuscript with input from all authors. G.S.K supervised the work.

## MATERIALS AND METHODS

### TCGA analysis

Publicly available transcriptomic data from breast cancer and normal human tissue obtained from the Cancer Genome Atlas (TCGA)^92^ were accessed via the UCSC Xena portal^93^. RNA sequencing data were used for the comparison of gene expression in normal (N), tumor (T) and metastatic tissues (MT) of Breast Invasive Carcinoma patients (N, N=292; T, N=1092; MT, N=7), via the TNMplot.com^94^. Sequencing reads were obtained by the Cancer Genomics Project^92^ using Illumina® platform. After quality control, expression data were normalized using RSEM (RNA-Seq by Expectation Maximization)^95^ and logged transformed (log2). For gene expression analysis and comparison between the breast cancer subtypes, the UALCAN web platform^96^ was used with the Cancer Genome Atlas (TCGA) RNA sequences (Normal, N=114; Luminal, N=566; HER2^+^, N=37; TNBC, N=116). Results were visualized as box and whisker plots using the ggplot2 R package (v3.2.1) and statistics performed with a Wilcoxon test applied to the comparison groups [Normal (N) vs Primary tumor (T) or metastatic (MT)] by using R version 3.5.1. For correlation analyses, RNAseq data were downloaded from xena- browser.net (ID: TCGA.BRCA.sampleMap/HiSeqV2, unit: log2(norm_count + 1), and corresponding phenotype data (ID: TCGA.BRCA.sampleMap/BRCA_clinicalMatrix). This dataset contained a total of 1247 samples, but only primary tumors were analyzed (n=1101). The R value from Pearson correlation between normalized RSEM counts of chemokines receptors/ligands and cell-type-specific genes was based on methodology developed by Tirosh et al. (2016)^97^, and was calculated using GraphPad Prism v10. Then, unsupervised hierarchical clustering gene tree was generated based on these correlations, using Multi Experiment Viewer (MeV) v4.9.

### Mice

All studies involving mice were carried out in accordance with the National Institutes of Health (NIH) regulation concerning the care and use of experimental animals. Procedures were approved by the Albert Einstein College of Medicine Animal Care and Use Committee. Transgenic mice expressing the Polyoma Middle T (PyMT) oncogene under the control of the mouse mammary tumor virus-long terminal repeat (MMTV-LTR), simply known as “MMTV-PyMT” or “PyMT” mice, have been previously documented to develop spontaneous mammary carcinomas and metastases^98^, and were bred in house.

PyMT mice were crossed with the Tie2^GFP^ reporter allele (JAX stock #003658)^99^, to generate double-transgene MMTV-PyMT; Tie2-GFP studies, used for single-cell transcriptomic studies. MAFIA mice [C57BL/6-Tg(CSF1R-EGFP-NGFR/FKBP1A/TNFRSF6)2Bck/J] were obtained from The Jackson Laboratory and were implanted with tumor pieces (2 mm × 2 mm) into the fourth mammary fat pad on the left side. Tumor cell injection xenografts were generated by orthotopic injection of 1x10^6^ MDA-MB-231 cells (shControl or shCXCR4-1 expressing) resuspended in sterile PBS with 20% type I collagen (Corning) into the right inguinal mammary fat pad of 8-week-old female mice with severe combined immunodeficiency (SCID; The Jackson Laboratory). Mice bearing xenografted tumors, either cell line-injected or patient-derived, were sacrificed as soon as tumors reached ∼0.5-0.7 cm in diameter.

### Single Cell Transcriptomics of the MMTV-PyMT; Tie2-GFP mouse model

*Sample processing and sequencing:* Tie2^GFP^ micro-dissected mammary tumors from PyMT-Tie2^GFP^ female mice were collected at 10 weeks. Tumor cells were dissociated using fine scissors and then proteolytic digestion was performed with the Collagenase Type IA and DNase buffer. Dissociated cells were then incubated for 30 minutes at 37°C with rotation followed by filtration using a 100 μM strainer. ACK buffer (Thermo Scientific) was used to lyse red blood cells and cells were subjected to dead cell exclusion using the MACS dead cell removal kit (Miltenyi). Tumor samples with viability greater than 70% were used for analyses. Isolated cells were processed for droplet based scRNA-seq using the 10X genomics Chromium Single Cell 3’ Library & Gel bead Kit V3.1 according to the manufacturer’s protocol. Sample inputs of 10,000 cells per lane were loaded onto each 10X channel. Libraries were sequenced on a Nextseq 2000 with 100 bp paired-end reads at the Einstein Epigenomics Core.

*Sc-RNA-seq quality control and pre-processing:* Cellranger v7.0.1 intron mode was used for alignment of reads to a custom mm10 mouse reference genome which included PyMT and GFP sequences. CellRanger was used to process and filter UMIs. Cells with less than 500 UMIs were filtered out for each sample. Pre-processing and filtration steps were applied to individual samples which were then concatenated together into a single dataset file. For doublet detection scvi-tools^100^ integrated SOLO^101^ algorithm was implemented. SOLO takes the pre-trained scvi-object as input and to obtain the same we first filtered the genes appearing in less than 10 cells followed by computing 2000 most highly variable genes (HVGs) by applying “flavor=seurat_v3” parameter. The filtered dataset was trained using scvi model and was used as input for SOLO algorithm. After doublet detection the dataset was re-loaded and predicted doublets were removed after iterative rounds and visualization on the clustered data. Further filtration steps include removing the cells that expressed less than 200 genes. QC matrix was implemented using scanpy^102^ to identify the percent of ribosomal and mitochondrial content in the samples. To further cleanup the data, we removed the cells that express unusually high number of genes.

The upper limit was detected using the 98^th^ percentile of n_genes_by_count and only retained the cells that expressed number of genes less than the upper limit. Cells were further filtered for less than 50% of the mitochondrial content. Ribosomal genes that start with ‘Rps’ and ’Rpl’ were removed from the samples. After the pre-processing steps, both the samples were concatenated using scanpy and we had 18188 number of cells in total from both the samples (8670 cells, Sample 1; 9518 cells, Sample 2).

*Data normalization and UMAP visualization:* Raw counts were normalized to the median library size and were log transformed in the scanpy. Post-normalization we selected 3000 HVGs using scanpy function of highly_variable_genes by applying “flavor=seurat_v3” parameter. We then performed the principal component analysis (PCA) using the log-transformed data of the selected HVGs and retained the first 30 PCs based on the percent variance. Scanpy neighbor function was used to compute UMAP^103^ embeddings on the PC representation and Euclidean distance and 30 nearest neighbors. The UMAP representation was then plotted using ‘min_dist=0.3’ using scanpy library function.

*Cell typing and gene expression signature:* To cluster cells into subgroups, we used leiden algo- rithm in scanpy tools function with resolution parameter set at 1 and k=30. To represent the phenotype similarities, we used PhenoGraph clustering with k=30 and resolution= 0.5. Next, we annotated the identified clusters in different cell types based on marker gene expression. We visualized the log-normalized expression of our target genes on the UMAP using scanpy plot function at vmax set between p99-p100.

### Single-Cell Transcriptomics of the NT2.5 Mouse Model

We recovered a previously published dataset^104^ including single-cell RNA sequencing on the HER2-overexpressing breast model, NT2.5, to examine the expression of the CXCL12/CXCR4 axis on tumor and stromal components. Specifically, for library preparation, 10× Genomics Chromium Single-Cell 3′ RNA-seq kits v2 were used. The gene expression libraries were prepared according to the manufacturer’s protocol. RNA was extracted from 20 tumors from the following experimental groups: Vehicle control (V), entinostat-treated (E), entinostat and anti-PD-1 (EP), entinostat and anti-CTLA-4 (EC), entinsotat with anti-PD-1 and anti-CTLA-4 (EPC), with four biological replicates from each of the five experimental groups. Tumors were sequenced in four batches: RunA (eight tumors; 1 E, 2 EP, 2 EC, 2 EPC, 1 V), RunB (eight tumors; 1 E, 2 EP, 2 EC, 2 EPC, 1 V), Pilot1 (two tumors, 1 E and 1 V), and Pilot2 (two tumors, 1 E and 1 V). Each batch had an approximately equal assortment of samples from each treatment group to reduce technical biases. Illumina HiSeqX Ten or NovaSeq were used to generate approximately 6.5 billion total reads). Paired-end reads were processed using CellRanger v3.0.2 and mapped to the mm10 transcriptome v1.2.0 by 10× Genomics with default settings. ScanPy v1.9.1 and Python v3 were used for quality control and basic filtering. For gene filtering, all genes expressed in less than 3 cells within a tumor were removed. Cells expressing less than 200 genes or more than 8,000 genes or having more than 15% mitochondrial gene expression were also removed. Gene expression was total-count normalized to 10,000 reads per cell and log transformed. Highly variable genes were identified using default ScanPy parameters, and the total counts per cell and the percent mitochondrial genes expressed were regressed out. Finally, gene expression was scaled to unit variance and values exceeding 10 standard deviations were removed. There were 54,636 cells and 19,606 genes after pre-processing. Batch effects were corrected using the ComBat batch correction package. Neighborhood graphs were constructed using 10 nearest neighbors and 30 principal components. Tumors were clustered together using Louvain clustering and 6 main clusters were identified (with resolution parameter 0.1).

### Inhibitors, compounds, and administration routes

AMD3100 has been extensively validated as a pharmacological suppressor of the CXCL12/CXCR4 pathway in certain contexts^23–27^. In this study, AMD31000 was administered at a dose of 5 mg/kg (diluted in PBS) by intraperitoneal (i.p.) injection, on a daily basis for two weeks. PBS alone was administered as vehicle-control. High molecular weight (155 kDa) tetramethylrhodamine (TMR)-dextran was administered at a dose of 100 µl of 10 mg/Kg (diluted in PBS) by tail vein (i.v.) injection, as described^29,38,105^, 1 hour before termination of experiments. For fixed-frozen immunofluorescence, the labeling of the flowing vasculature has been described^29^, by administering anti-mouse CD31-biotin by tail vein injection (i.v.), 10 minutes before the termination of the experiments. For macrophage depletion with B/B Homodimerizer in MAFIA Mice, animals were administered 10 mg/kg B/B homodimerizer (AP20187; Clontech) diluted in 4% ethanol, 10% PEG-400, and 1.7% Tween-20 or vehicle control by intraperitoneal injection daily for 5 days.

### Transduction of MDA-MB-231 cells using lentiviral vectors

The human embryonic kidney cell line, HEK293T, and the human breast cancer cell line, MDA-MB-231, were cultured in Dulbecco’s Modified Eagle Medium (DMEM; ThermoFisher) supplemented with 10% fetal bovine serum (FBS) and antibiotic (penicillin/streptomycin, 1%). MDA-MB-231 cells were used between passages 20-25 and HEK293T cells were used between passages 5-10. Cells were grown in a humidified 5% CO_2_ incubator at 37°C and routinely shown to be mycoplasma free. Short hairpin RNA (shRNA) backbone vectors specific against human CXCR4 (#SHCLNG-NM001008540.2; shCXCR4-1, TRCN0000004052; shCXCR4-2, TRCN0000004053; shCXCR4-3, TRCN0000004054; shCXCR4-4, TRCN0000004055) and the TRC empty vector (Control, SHC001) were purchased from Einstein’s Molecular Cytogenetic Core. Lentiviral particles were generated by transfecting confluent HEK293T cells with a 1.25 μg backbone shRNA plasmid and 3.75 μg VIRAPOWER lentiviral expression system (Invitrogen) using 48 μg of Transporter 5 Transfection Reagent (Polysciences), following manufacturer’s instructions in T25 flasks. The viral supernatants were harvested after 72 hours, centrifuged at 1000xg for 10 minutes and filtered through a 0.45 μm filter. T25 flasks of MDA-MB-231 cells at 25-30% confluency, were transduced with 2 ml viral supernatant, 3 ml complete growth medium, and 12 μg/ml polybrene (Sigma) for 5 hours. The stable transductants were identified by maintaining the cells under puromycin selection (1μg/ml) for 2 weeks.

To confirm knockdown of CXCR4 in the stable cell lines, whole cell extracts were prepared from the stably selected shControl and shCXCR4 MDA-MB-231 cell lines by resuspending cell pellets in 1.5× Laemmli sample buffer containing 2% 2-mercaptoethanol and incubating at 95°C for 10 min before storing at −20°C. Proteins were separated on 10% sodium dodecyl sulfate polyacrylamide (SDS-PAGE) gels and transferred to nitrocellulose blotting membranes (Amersham Protran 0.2μm). The membranes were probed with antibodies directed to CXCR4 (1:1,000; BD Pharmingen) and GAPDH (1:10,000; Abcam) overnight at 4°C. The membranes were washed three times for 5 min using 0.1% Tween-20 in Tris-buffered saline (TBS) before incubating for 1 hour with secondary antibodies (Li-Cor IRDye 680RD) directed against the appropriate species. Following three 5-min washes with 0.1% Tween-20 in TBS, membranes were imaged on a Li-Cor Scanner and processed with ImageJ.

### Circulating tumor cell assay

Circulating tumor cells (CTCs) were collected from the left atrium of the heart as terminal procedure, and quantified as previously described^105^. The values were first normalized per mL of blood collected, and then expressed as fold-difference to the mean of the control group.

### In vivo invasion assay

Cell collection in microneedles placed in the primary tumor of MMTV-PyMT mice was carried out as previously described^31^. Briefly, mice were anesthetized using isoflurane, and invasive cells were collected in 33-guage Hamilton needles, filled with Matrigel (Beckton Dickinson) containing 62.5 nM CXCL12 (R&D systems), or PBS alone, for 4 hours^20^. At the conclusion of tumor cell collection, the contents of the Hamilton needles were extruded with PBS containing DAPI onto a glass coverslip. The number of cells in each needle was counted, and then normalized to the control samples for each mouse.

### TMEM doorway immunohistochemistry

TMEM doorway immunostaining and quantification in mice were performed using a previously described method of triple-stain immunohistochemistry (IHC) for macrophages (IBA1; brown), endothelial cells (EMCN; blue), and tumor cells (MENA; red)^105^. The “TMEM doorway score” represents the total number of TMEM doorways in 10 high power fields (HPFs) of murine tumor tissue, and has been calculated by an experienced pathologist (M.H.O)^35,37^.

### Multiplexed immunohistochemical consecutive staining on single slide (MICSSS)

We collected invasive ductal carcinomas from 6 patients with hormone receptor positive/ HER2^-^ breast cancer that were treated with adjuvant systemic therapy (anti-estrogen therapy, with or without chemotherapy). 3 patients were of white and 3 of Black ancestry. Within each racial group, 2 patients developed distant recurrence and 1 did not. Formalin-fixed paraffin-embedded (FFPE) tissue slides from these patients were subjected to multiplexed immunohistochemical consecutive staining on a single slide (MICSSS)^50^. An automated immunostainer (Leica Bond RX, Leica Biosystems) was used to bake slides containing human invasive ductal carcinoma and perform iterative staining, consisting of chromogenic revelation via 3-amino-9-ethylcarbazole (Vector Laboratories) and counterstaining with hematoxylin as described before^106^. After each round of staining, slides were removed, mounted with a glycerol-based mounting medium, and scanned to obtain digital images using an Aperio AT2 scanner and Aperio ImageScope DX visualizer software v.12.3.3 (Leica). After scanning, slide coverslips were removed in hot water (∼50 °C) and tissue sections were bleached. This process was repeated for the length of an optimized 10-marker panel consisting of PD1, FoxP3, CD8, PD-L1, CD206, PanCK, CD68, CD31, Granzyme B, and CD20, as described before^106^.

After all the scans were acquired, the images were aligned down to the single cell level. To align images, a project was created in QuPath (v0.4.3) and all digital whole slides were imported. Next, the QuPath project file was opened in Fiji using the Big Data Viewer plugin, and the PD1 image was selected as the “fixed reference” image, while the other nine images were designated as moving sources. QuPath’s "Create Warpy Registration" tool was used with default parameters, and rigid and affine registrations were sequentially performed to align the images. Next, the rectangle ROI tool was used to draw an ROI surrounding the tissue of interest, and the landmark grid spacing was changed to 800 microns. After processing, a JSON transformation file was generated. This alignment procedure was repeated for each scan (as moving source), keeping the source image (PD1), as the fixed reference source. Finally, the different JSON transformation files were used in QuPath’s “Interactive Image Combiner Warpy” tool to create a multiple image overly consisting of all ten aligned scans.

For detection of cell subsets using one marker, QuPath’s cell detection function was used to identify individual cells within regions-of interest. This detection consisted first of selecting the annotations for either “Intravasation Portal” or “non-Intravasation Portal” regions. Next, the cell nuclei within the selected region was detected based upon the Hematoxylin stain. Finally, the areas corresponding to cell nuclei were expanded by 1.5 µm, to demarcate the total area of each cell. Briefly, this procedure consisted of selecting the Cell Detection module in QuPath, selecting the PD1-Hematoxylin image as the cell detection channel, and setting the pixel size to 1 micron, detection threshold to 0.1, and expansion criteria to 1.5 microns, and the “include cell nuclei” option to “disabled”. All other options were kept at default settings. Once cells were detected, their positivity for the different markers could be assessed using the Create Single Measurement Classifier module. After selecting the DAB channel for the marker of interest, the object filter was set to “Detections (all)”, the Measurement parameter was set to the mean intensity of the cell, and the Above Threshold selector was set to the marker of interest. The thresholds for detection were determined for each channel manually above the level of the background. The threshold for positivity was then finetuned by using the preview of the cells selected for each threshold chosen (i.e. enabling the Live Preview checkbox) and adjusting the threshold slider to reduce the number of non-specific cells identified. Once all settings were configured, the classifier was saved as a file, and the settings applied to the image. For detection of cell subsets using two markers, were identified by using the Create Composite Classifier module and selecting the two previously identified single positive cell classifiers. The results for all individual cells were exported as a CSV file.

### Multiplex immunofluorescence

Tumors or part of the tumors were removed at the time of sacrifice and fixed overnight in 4% PFA, cryoprotected in 30% sucrose and then frozen in OCT (for fixed frozen IF) or fixed for 48 h in 10% formalin and then embedded in paraffin (for FFPE IF). In all instances, 5um- thick sections were cut, and staining was performed as described, using either conventional IF for fixed-frozen samples, or Tyramide Signal Amplification (TSA) IF for FFPE tissues ^28,29,105^. The following primary antibodies were either used alone or multiplexed in various combinations: goat anti-Mrc1/CD206 (R&D Systems), rat anti-CXCR4 (BD Biosciences), rat anti-ZO1 (clone R40.76; Millipore), rat anti-EMCN (clone V.7C7; Santa Cruz Biotechnology), rabbit anti-CD105 (clone; abcam) mouse anti- CXCL12 (clone K15C; Millipore), rabbit anti-IBA1 (Wako), rabbit anti-CD31 (Cell Signaling), anti-TMR (A-6397; Life Technologies), mouse anti-MENA (BD Biosciences), rat anti-B220/CD45R (MAB1217; R&D), rabbit anti-PDGFRβ (3169S; Cell signaling), rabbit anti-CD4 (ab183685; abcam), rabbit anti-CD8 (ab217344; abcam), mouse anti-PANCK (C2562; Sigma-Aldrich), rat anti-PyMT (MA1-46061; Invitrogen), and chicken anti-MENA^INV^ (in-house generated^105^). Nuclei were stained with DAPI, and samples were mounted with Prolong Gold antifade reagent (Molecular Probes/Invitrogen). Tissue slides were imaged on a 3DHISTECH P250 Flash III digital whole slide scanner, using a 20x 0.75NA objective lens. Tissue suitable for scanning was automatically detected using intensity thresholding. Whole-tissue images were uploaded in CaseViewer version 2.2 (3DHISTECH), and regions of interest (ROIs) were captured with the Image Snapshot tool, as TIFF images. All quantitative analyses were performed on raw 8-bit TIFF images in ImageJ or QuPath.

*TMEM doorway activity:* The activity of TMEM doorways is assessed via the calculation of TMEM doorway-mediated vascular permeability, a prior developed assay^38,105^ that measures the extravasation of 155-kDa dextran, conjugated to tetramethylrhodamine (TMR) into the tumor tissue. The Assessment of tight junction integrity at TMEM doorways was performed via the quantification of Zona Occludens-1 (ZO1) protein, coupled to a modified extravascular dextran analysis, using multiplex IF in fixed-frozen tumor samples, as previously described^29^.

*CXCL12 expression gradient:* To measure CXCL12 expression gradient around TMEM doorways, the triple-IHC TMEM stain (used to identify TMEM doorways) was co-registered with a sequential IF section stained with Endomucin (EMCN) and CXCL12. TMEM doorways were identified by a pathologist (M.H.O.) independently of the co-registration analysis, to avoid any operator-based biases for TMEM doorway selection based on CXCL12 expression levels. To measure CXCL12 expression gradients around the perivascular niches (PNs), the IF section, co-stained with anti-EMCN and anti- CXCL12, was directly used to generate a blood vessel mask, and was enlarged by 10µm, corresponding to the mean size of a tumor cell in the PyMT model, creating the PN ROI. In both cases, the “Enlarge” feature of ImageJ was used to designate concentric annuli (with 3µm thickness) around TMEM and PN ROIs. The “Fill Holes” feature of ImageJ was also used in the thresholded, binarized EMCN channel. The resulting EMCN ROIs were then applied as exclusion masks to the TMEM- and PN-generated annuli so as to avoid including the vascular lumen when measuring chemokine expression in the tumor microenvironment. Finally, CXCL12 was thresholded above the secondary antibody (negative) control level, binarized and quantified in each ROI annulus as a percentage area with signal above the threshold. Data were presented as ratios of the cumulative CXCL12^+^ area divided by the total ROI area for each additional annulus away from TMEM or PN ROIs. To further ensure specificity, we additionally employed an “absorption” control in our analysis to limit experimental noise originating from the non-specific reactivity of the mouse anti-CXCL12 antibody with mouse IgG present within the tissue. Tissue sections were pre-incubated with a rabbit anti-CXCL12 antibody (clone D32F9; Cell Signaling) for 30 mins, followed by standard incubation with primary mouse anti-CXCL12 antibody and amplification with secondary anti-mouse IgG antibody for signal detection. Indeed, we confirmed non-specific reactivity of the mouse IgG with the anti-CXCL12 antibody (data not shown). However, the non-specific signal was reduced during the thresholding step in the pipeline described above.

*Quantification of CXCR4^+^MENA^INV+^ cells in 45µm radius-long ROIs:* To measure CXCR4^+^MENA^INV+^ tumor cells around TMEM doorways, the triple-IHC TMEM stain was first used to identify TMEM doorways, and the slide was then aligned and co-registered with a sequential IF section stained with EMCN, MENA^INV^ and CXCR4. Based on findings from TMEM-generated CXCL12 chemokine gradients, a circular ROI of 45µm radius was drawn around each TMEM doorway with the corresponding TMEM doorway drawn as a triangle (triad of cells composing the TMEM doorway) by the pathologist and positioned at the center of this circle. The number of CXCR4^+^MENA^INV+^ tumor cells was quantified and presented as a percentage among all DAPI^+^ cells within the 45µm-radius TMEM ROIs. Numbers of CXCR4^+^MENA^INV+^ cells were also quantified in the entire fields of view for comparison, not only in the TMEM ROIs.

*Disseminated tumor cells (DTCs) in bone marrow and lungs:* To detect single disseminated tumor cells within the parenchyma of bone marrow and lungs, multichannel immunofluorescence staining for PANCK and PyMT was performed, along with EMCN, to define vascular architecture of these tissues. The analysis was conducted exclusively in Caseviewer v2.2. For each case, approximately 5 mm^2^ of bone marrow and lung tissue parenchyma was captured as an ROI using the polygon tool. DTCs were then manually quantified within these ROIs on the basis of double PANCK/PyMT positivity. The density of DTCs in each secondary metastatic site was expressed as number of DTCs per mm^2^ of lung or bone marrow parenchyma.

*Quantification of CD8^+^ cells in 45µm radius-long ROIs:* Similarly, to measure CD8^+^ T cells around TMEM doorways, the triple-IHC TMEM stain was also used to identify TMEM doorways, and the slide was then aligned and co-registered with a sequential IF section stained with EMCN and CD8. Based on findings from TMEM doorway-generated CXCL12 chemokine gradients, a circular ROI of 45µm radius was also applied. The number of CD8^+^ T cells was quantified in 10 low power fields of view (LPF, 20x magnification), and presented as an average absolute number of the total CD8^+^ T cells per LPF. Then the number of CD8^+^ T cells was quantified within these 45µm-radius TMEM ROIs and presented as an average absolute number of the CD8^+^ T cells within the 45µm-radius TMEM ROIs per LPF, which was finally normalized to the total CD8^+^ T cells per LPF.

*Homogeneity analysis of CD8^+^ T cell distribution:* We developed a new morphometric feature to characterize the spatial distribution of CD8^+^ T cells. Up to five representative, low resolution 10x FOVs depicting the tumor area were captured using CaseViewer v2.4 (3D HISTECH), and uploaded on QuPath v0.5.1. The “DAPI ROI” was automatically demarcated by a trained pixel classifier using these snapshots, to distinguish tissue-empty areas and areas corresponding to dead cells or necrosis. The CD8^+^ T cell population was also automatically demarcated and counted, using the cell detection tool with the following settings: requested pixel size 0.5 μm; background radius 8 μm, sigma 5 μm; Min/Max Area 60-400 μm^2^; and threshold 10. Slight modifications to these settings were made across different batches of staining, depending upon the quality/pattern of staining. The spatial arrangement of CD8^+^ T cells in the tumor parenchyma, here termed “homogeneity” (h) of distribution, was calculated. A series of modifications on the already obtained FOVs has been performed, immediately following the identification of the “DAPI ROI” using the pixel classifier, and counting of the cells of interest using the cell detection tool, as described above. First, a “template ROI” was applied to the image, to segment the “DAPI ROI” into 20 equally-sized, rectangular-shaped sectors (i). Because up to 5 fields-of-view per animal were retrieved, the total number of sectors varied between 20 and 100 per animal. The density of CD8^+^ T cells was then independently calculated for each sector *i*. The standard deviation of all the sector densities of CD8^+^ T cells was then calculated for a reference group (SD^ref^), across all fields-of-view. We selected the control group (i.e., vehicle-treated) as the reference group, because we assumed that any deviations observed within the control group should possess certain physiological/biological relevance. Then, each sector was subjected to the following logical check, independently, and the number N of TRUE returns was counted for each mouse:

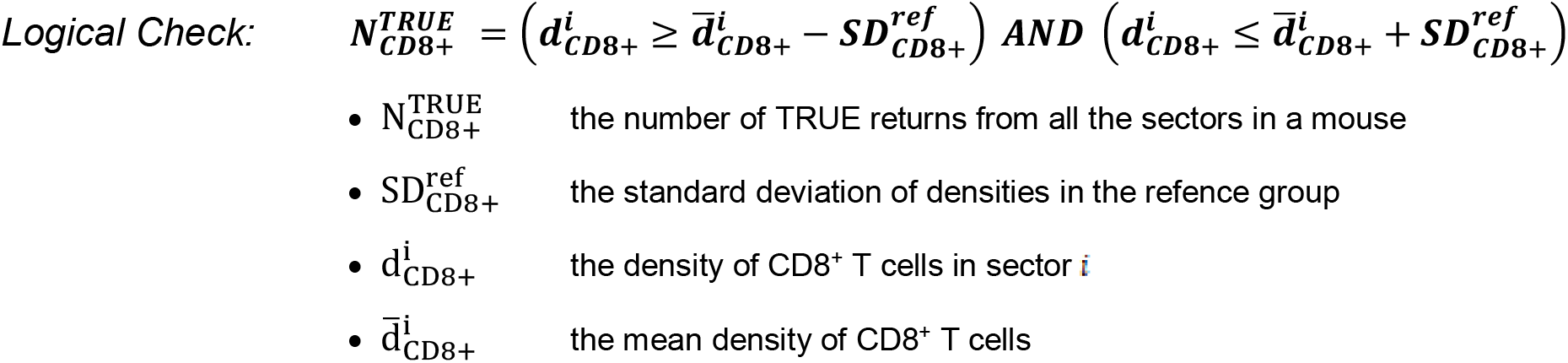

This logical check returns TRUE if the sector’s density falls within 1 SD^ref^ from the mean density, and FALSE otherwise. Following these determinations, the homogeneity index, i.e., h_CD8+_, reports the fraction of sectors that checked TRUE on these logical checks, for each subset of interest. The homogeneity index shows the fraction of sectors whose density distribution falls within an expected variance (i.e., standard deviation of a reference group). A high h_CD8+_ index informs that most sectors tend to have similar densities of CD8^+^ T cell subsets. A low h_CD8+_ index, however, informs that a large number of tumor sectors presents with either very-high or very-low density of CD8^+^ T cells, thus implying that the overall spatial distribution/arrangement within the tumor is more heterogeneous and clustered.

*Vasculature analyses:* To measure angiogenic parameters, we co-stained mammary tumor slides with DAPI, CD105 and EMCN. After scanning, 5 snapshots at 20x magnification were taken as TIFF, and each channel was saved independently. For Microvessel density, EMCN channels were subjected to the following pipeline: After being transformed into 8-bit in Fiji, the images were thresholded (default, auto), creating binary masks, which were then processed using the commands “Despeckle”, “Remove Outliers” and “Fill holes” in a sequence. The resulting masks were quantified, and expressed as area percentage (%) of the entire field. Additionally, we conducted skeleton analysis for in-depth analysis of the topological traits of the vasculature network. The calculated features were: (a) number of branches, defined as the sum of the branches of all the EMCN FOV_S_ for each animal; (b) Number of Junctions, defined as the sum of the junctions of all the EMCN FOVs for each animal; and (c) Average Branch Length, defined as the average of all branch lengths >=0.5 of all the FOVs for each animal. Finally, we calculated the CD105^+^ fraction within the entire vasculature network, to determine neoangiogenic and putatively “active vascularization”^42^. This fraction was calculated, first, by binarizing the CD105 signal within the EMCN mask in each FOV using intensity thresholding, and second, by applying the AND logic gate to demarcate the CD105^+^EMCN^+^ mask as an independent ROI. The surface area of this ROI was expressed as a fraction of the total EMCN ROI, calculated for Microvessel density, and as such, expressed as a fraction of tumor vasculature.

*Analysis of endothelial tight junction integrity:* Quantification of relative expression of blood vessel marker CD31, tight junction protein ZO1 and Dextran-TMR, was performed in fixed-frozen tissues using multiplex IF, as previously described^29^. All the individual channels were first thresholded to the level just above the background, as described^29^. For each FOV, the total TMR^+^ and ZO1^+^ areas were normalized to the respective CD31^+^ area, and finally expressed as a mean of all FOVs for each mouse.

### Flow Cytometry

Tumors were dissected and minced into 2-4 mm fragments, which were incubated in HBSS medium with 10 UI/mL collagenase I, 400 UI/mL collagenase D and 40 µg/mL DNase at 37℃ for 1 hr under agitation. After the samples were digested in the enzymatic mix, the fragments were homogenized, filtered through a 100µm nylon mesh cell strainer and cells were centrifuged at 1,450 rpm for 10 min at 4℃. Cell suspensions were incubated with 2.4G2 Fc Block Ab and stained in PBS 1%FCS, 2mM EDTA, 0.02% sodium azide with fluorescently tagged Abs purchased from eBioscience, BD Biosciences, R&D systems, or BioLegend (CD45, CXCR3, CXCR4, Ly6G, Ly6C, CD11c, KLRG1 CD103, PD-1 (CD279), MHCII (I-A/I-E), CTLA-4, PD-L2 (CD273), B220 (CD45R), Ki67, F4/80, CX3CR1, CD62L, TIGIT, TIM-3 (CD366), CD8a, CD3, NK1.1, CD169 (siglec1), CD11b, CD4, CD19, PD-L1 (CD274), Granzyme B, CD86, FOXP3). Brilliant stain buffer (BD) was used when more than two Abs were conjugated with BD Horizon Brilliant fluorescent polymer dyes. Cells were stained for cell-surface marker expression, then fixed in eBioscience Fixation/Permeabilization buffer prior to intracellular granzyme B and Foxp3 transcription factor (TF) staining in eBioscience Permeabilization buffer for 1hr. Data acquisition was done using a Cytek Aurora flow cytometer. All flow cytometry data were analyzed using FlowJo v9 or v10 software (TreeStar).

### Statistical analysis

Individual animals in each experiment are presented as points on dot plots. The horizontal lines indicate means and error bars represent standard deviations or standard errors of the mean. All *in vivo* experiments were performed using three independent animal cohorts, which were pooled together for the final presentation of the data. Using sample size calculations, we ensured that our studies have a probability of type II error of less than 20% (i.e., the statistical power is >80%). Statistical significance was determined by Mann Whitney U-test for two-group comparisons, and repeated-measures ANOVA for CXCL12 gradient expression analyses. Statistical significance was determined at the 0.05 level. Graphing and statistical hypothesis testing were performed in GraphPad Prism 8.2.1.

