## Supplementary figures for "Intratumoral CXCL12 Gradients Contextualize Tumor Cell Invasion, Migration and Immune Suppression in Breast Cancer"

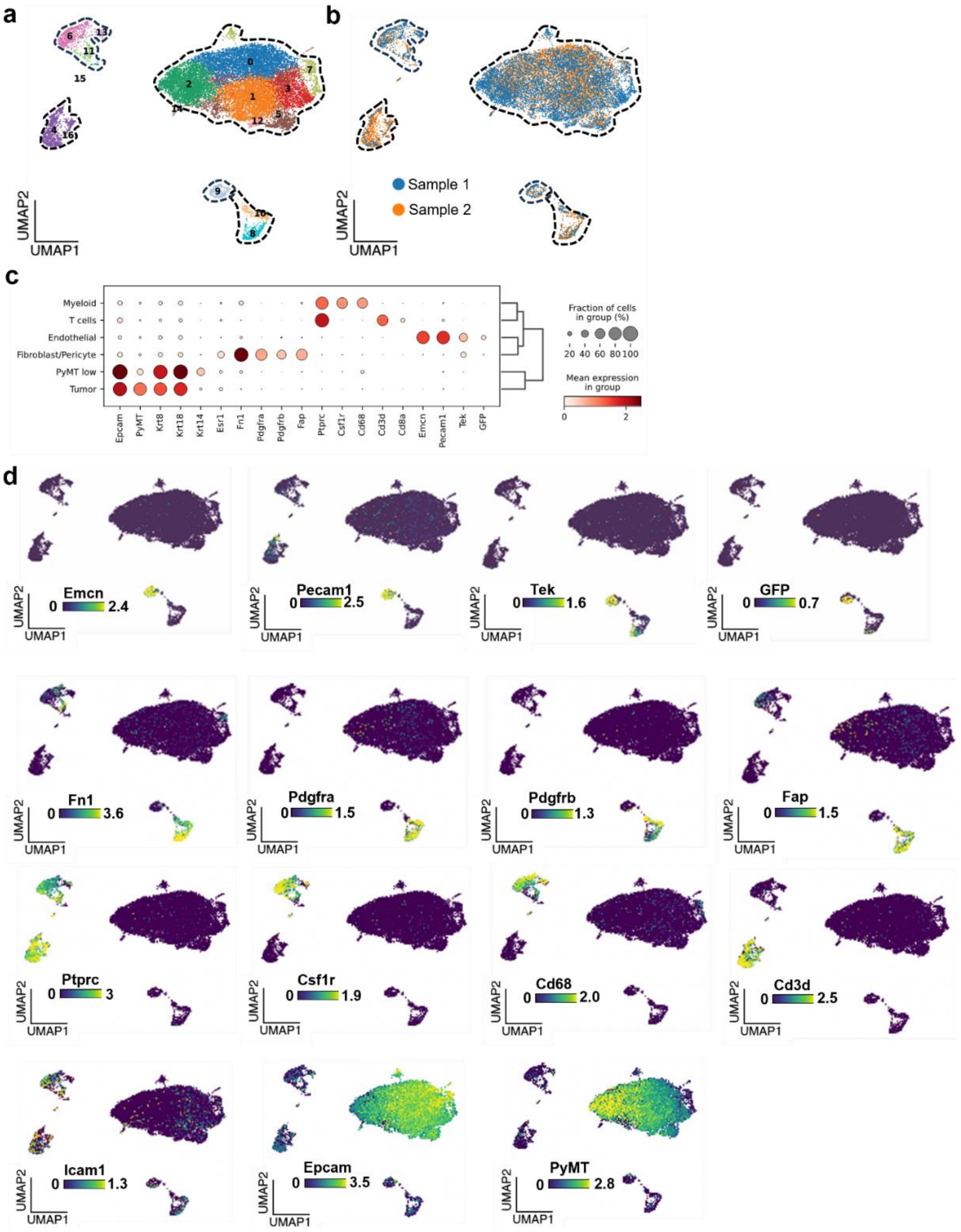


**Supplementary Figure 1. Single-Cell RNA Sequencing (scRNA-seq) of the MMTV-PyMT; Tie2-GFP mouse model of breast carcinoma. (a)** UMAP visualization of PyMT carcinomas profiled by scRNA-seq, indicating the extended clusters of cells with similar gene expression profile. **(b)** UMAP visualization of PyMT carcinomas profiled by scRNA-seq, indicating the distribution of cell populations originating from different samples analyzed. **(c)** Expression profiles of cell clusters for the indicated signature genes. The fractions of each cluster expressing a particular gene, and their expression levels are depicted according to the scales shown on the right. Dot color represents the z-score of the mean expression levels of the gene in the respective cluster, and dot size represents the fraction of cells in the cluster expressing the gene; gene names used for cluster definition are shown on the x axis. **(d)** expression profiles of the indicated gene signatures.

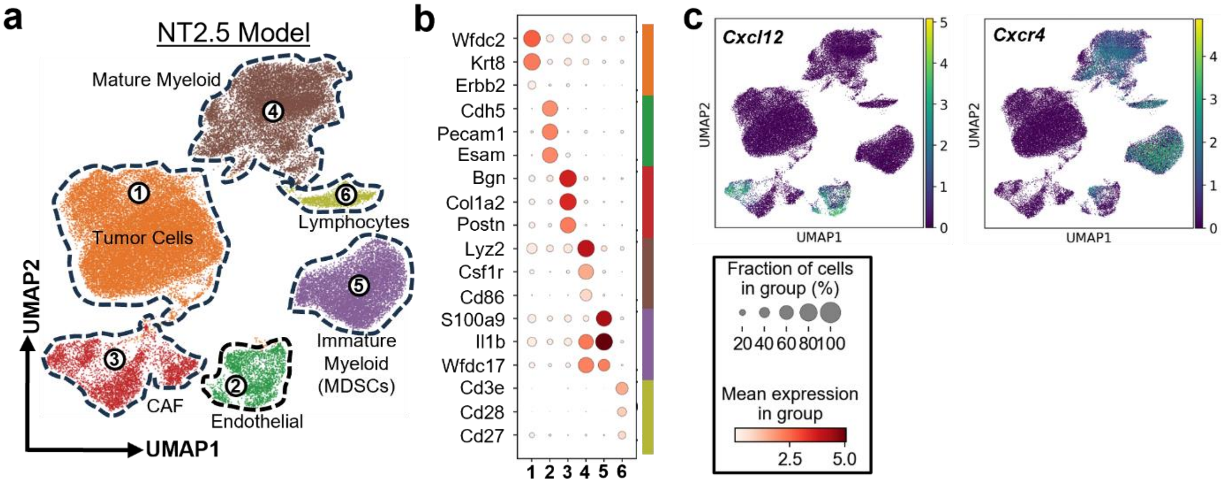


**Supplementary Figure 2. Single-Cell RNA Sequencing (scRNA-seq) of the NT2.5 mouse model of breast carcinoma. (a)** UMAP visualization of NT2.5 carcinomas profiled by scRNA-seq, indicating the cell clusters with similar gene expression profile. **(b)** Expression profiles of cell clusters for the indicated signature genes. The fractions of each cluster expressing a particular gene, and their expression levels are depicted according to the scales shown. Dot color represents the z-score of the mean expression levels of the gene in the respective cluster, and dot size represents the fraction of cells in the cluster expressing the gene; gene names used for cluster definition are shown on the x axis. **(c)** Expression profiles of *Cxcl12* and *Cxcr4*, two genes of interest in the current study.

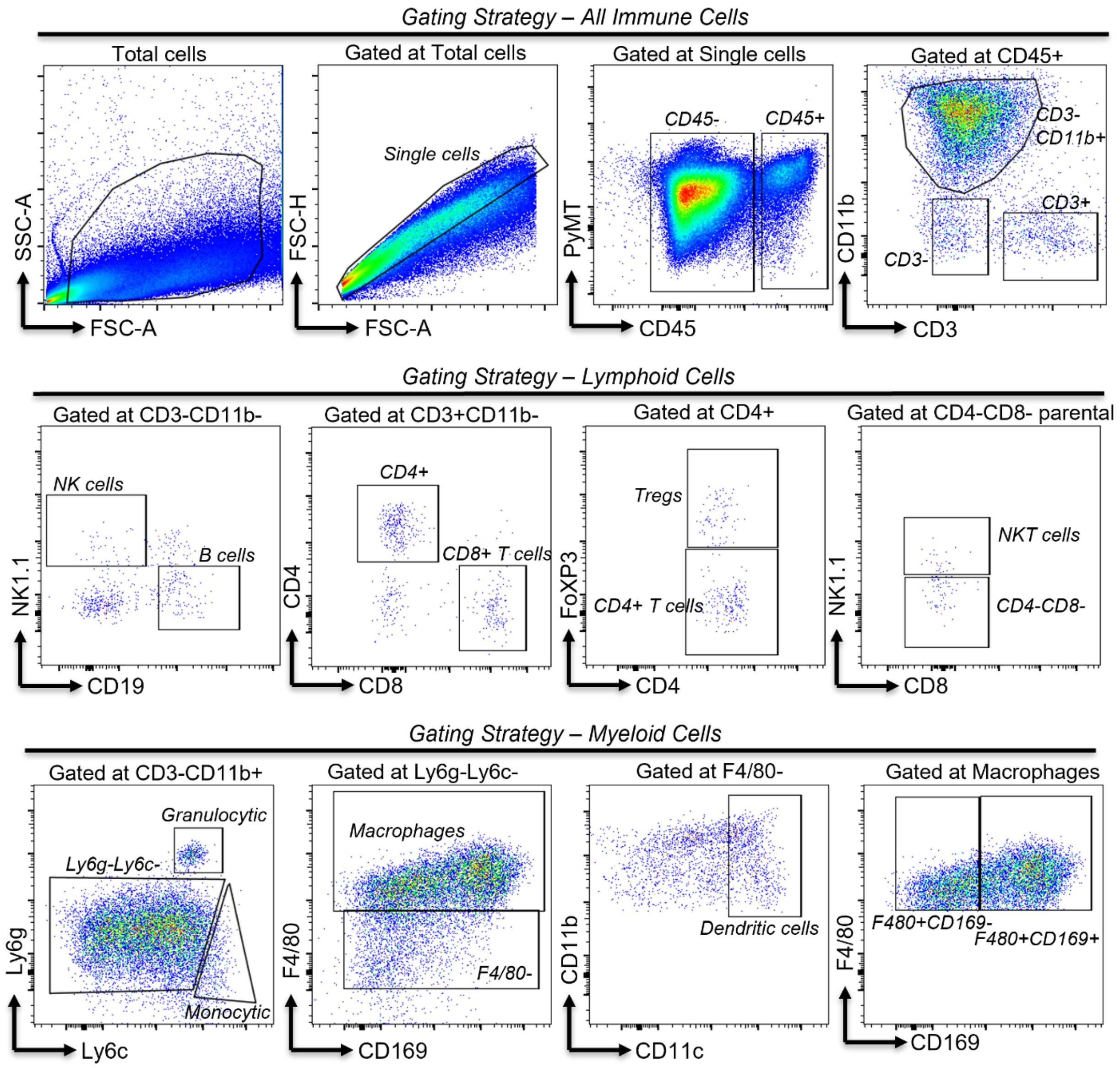


**Supplementary Figure 3. Overview of flow cytometry gating strategy utilized, comprised of multiple lineage markers to assess the tumor immune microenvironment.** The sequential gating strategy is illustrated in the flow cytometry plots. Starting from the initial singlet cell discrimination using forward scatter (FSC) and side scatter (SSC) parameters, immune cells were initially selected on the basis of increased CD45 expression, and subsequently gated to identify specific immune cell populations, using the described lineage markers.

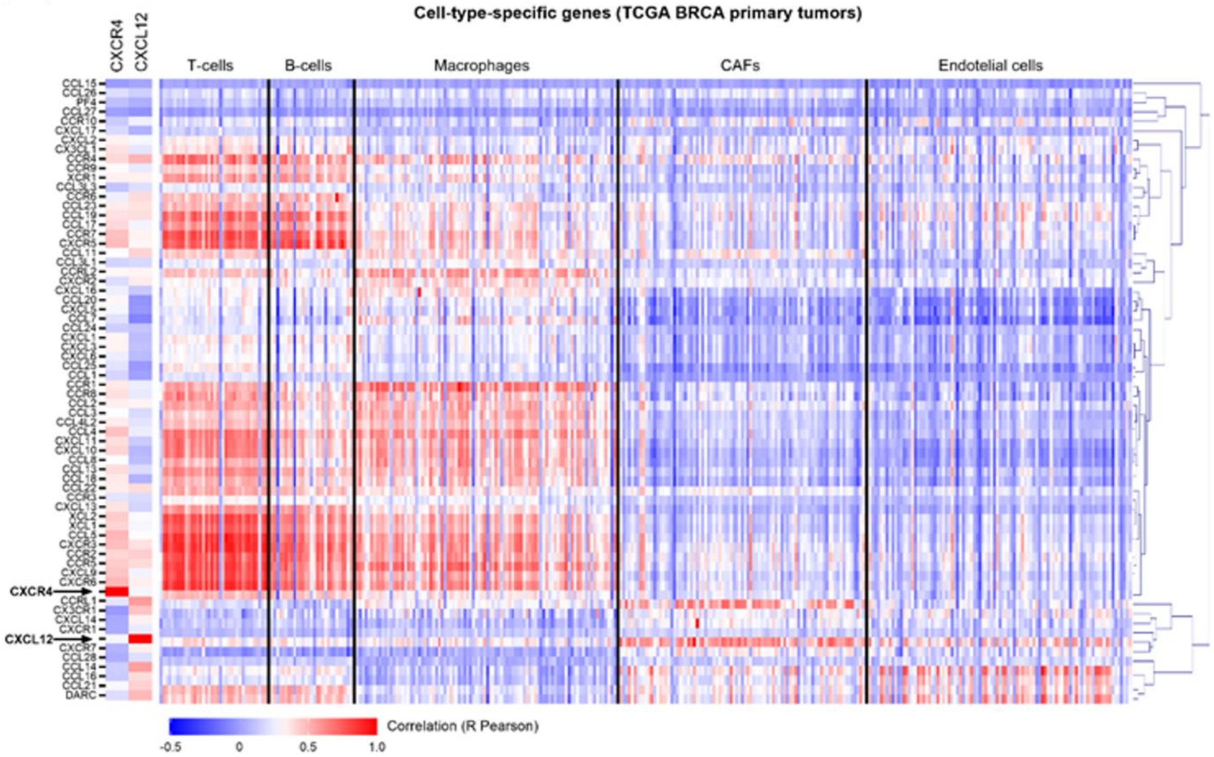


**Supplementary Figure 4. An extended hierarchical-cluster heatmap revealing correlations between all the chemokine receptor and ligand gene expression and different cell types in all the tumors.** The cell types [T cells, B cells, Macrophages, Cancer-Associated Fibroblasts (CAFs) and Endothelial cells] are defined by unique sets of cell-type-specific genes (See Supplementary Table 1)

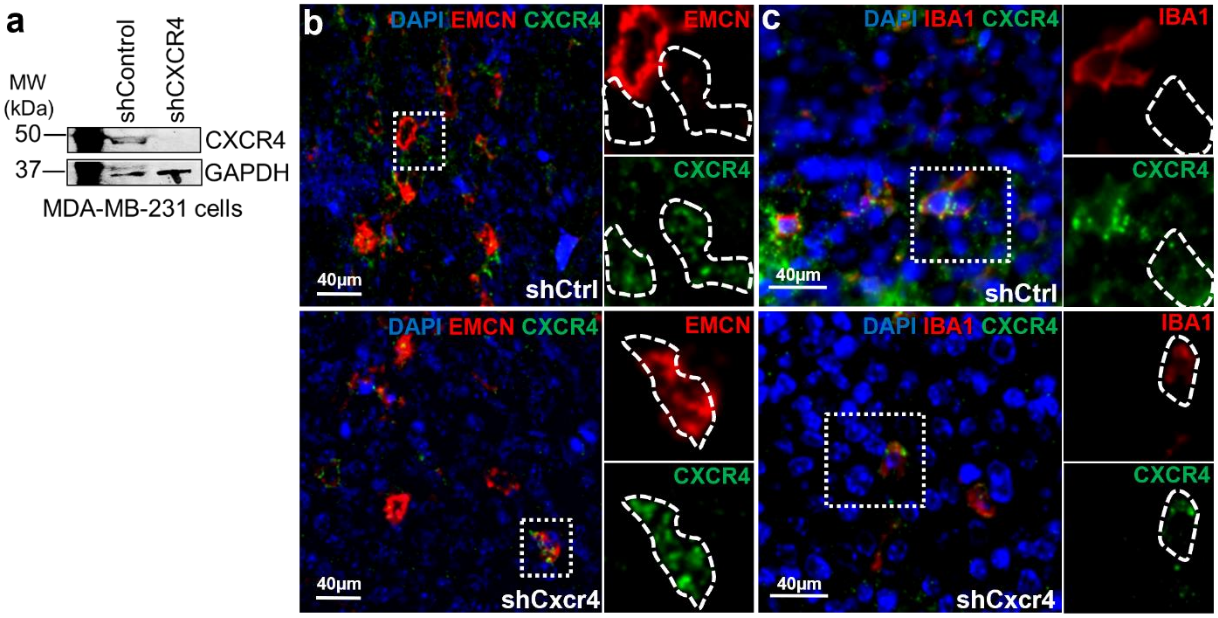


**Supplementary Figure 5. Development of MDA-MB-231 subline and mouse xenografts lacking expression of CXCR4 receptor. (a)** Expression of CXCR4 in MDA-MB-231 cells induced with either shControl or shCXCR4, revealing minimal or absent expression of CXCR4, in vitro. **(b-c)** Following injection of 231-shCxcr4 cells into SCID mice, the *in vivo* expression of CXCR4 is lost from tumor cells, but not from $CXCR4^+$ populations in the host mouse such as $EMCN^+$ blood vessels, and $IBA1^+$ myeloid cells, as indicated by multichannel immunofluorescence (lower rows in b and c). However, in mice developing 231-shCtrl tumors, there exist $CXCR4^+$ tumor cells (outlines) adjacent to $CXCR4^+$ blood vessels and macrophage (magnified inserts).

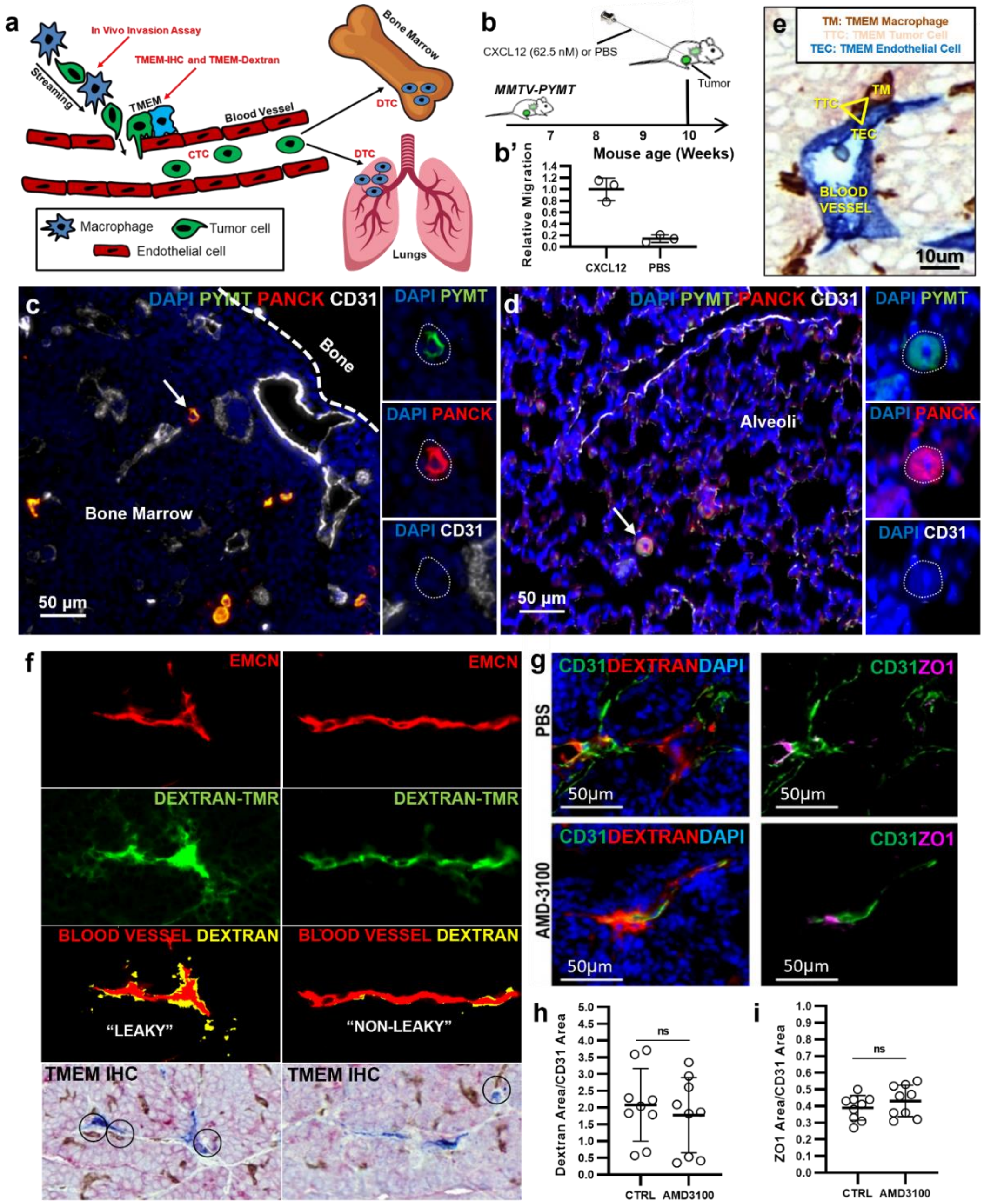


**Supplementary Figure 6. Assays for the assessment of cancer cell dissemination. (a)** An illustration depicting the cellular prerequisites of cancer cell dissemination, i.e., streaming cancer cell population, and TMEM doorway formation, along with key assays (written in red) for the analysis of cancer cell dissemination. **(b)** *in vivo* assay setup and **(b')** a control experiment demonstrating minimal or absent passive collection of tumor cells into Hamilton Needles, when they are coated with PBS, instead of a chemokine, in this case CXCL12. **(c-d)** Disseminated tumor cells (DTCs) in the bone marrow (c) or lung (d) are defined as PyMT$^+$ and Pancytokeratin$^+$ (PANCK$^+$) using multichannel immunofluorescence. **(e)** TMEM doorway triple-immunohistochemistry. **(f)** TMEM doorway activity assessment, by quantifying extra-vascularly leaked dextran-tetramethylrhodamine, using multichannel immunofluorescence. A leaky (1$^{st}$ column) and non-leaky (2$^{nd}$ column) blood vessel profile is seen. **(g-i)** Assessment of extravascular dextran leakage 1$^{st}$ column and quantification in h) and tight junction integrity (i.e., ZO1 expression; 2$^{nd}$ column and quantification in i), using multichannel immunofluorescence in MMTV-PyMT mice treated with either vehicle (PBS) or Amd3100.

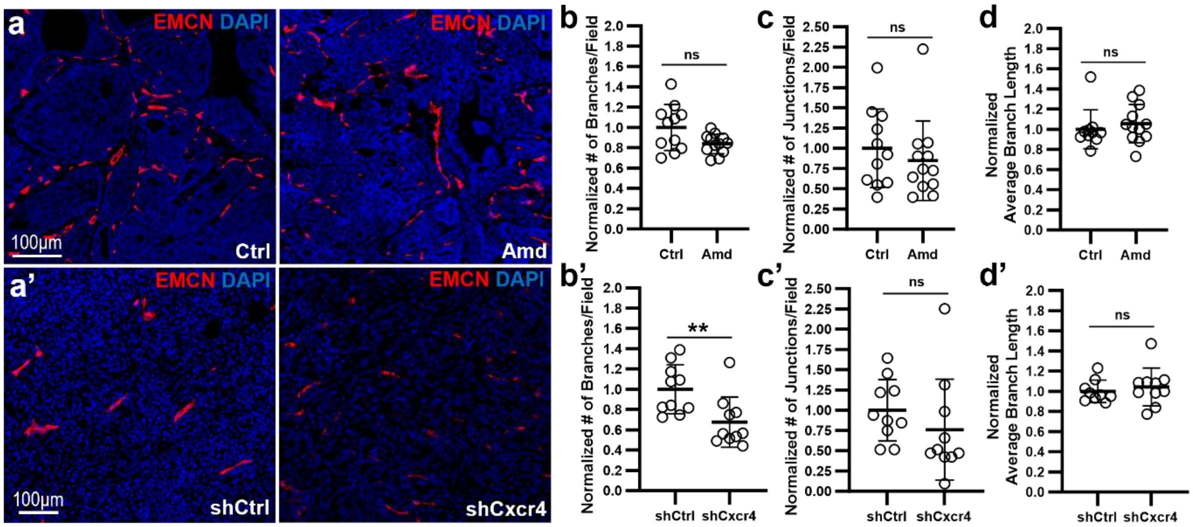


**Supplementary Figure 7. Assessment of branching morphogenesis in intratumoral blood vessels, using digital morphometry and skeleton analysis. (a-a')** Representative images of endomucin (EMCN) staining from PyMT mice receiving vehicle or Amd3100 (a), or mouse xenografts receiving 231-shCtrl or -shCxcr4 cell injections. **(b-b')** Quantification and normalization of the number of blood vessel branches per high-power field in the mouse models described in (a-a). **(c-c')** Quantification and normalization of the number of blood vessel junctions per high-power field in the mouse models described in (a-a'). **(d-d')** Quantification and normalization of average branch length in the mouse models described in (a-a'). *Mann-Whitney U-test; ns, non-significant; ** p<0.01.*

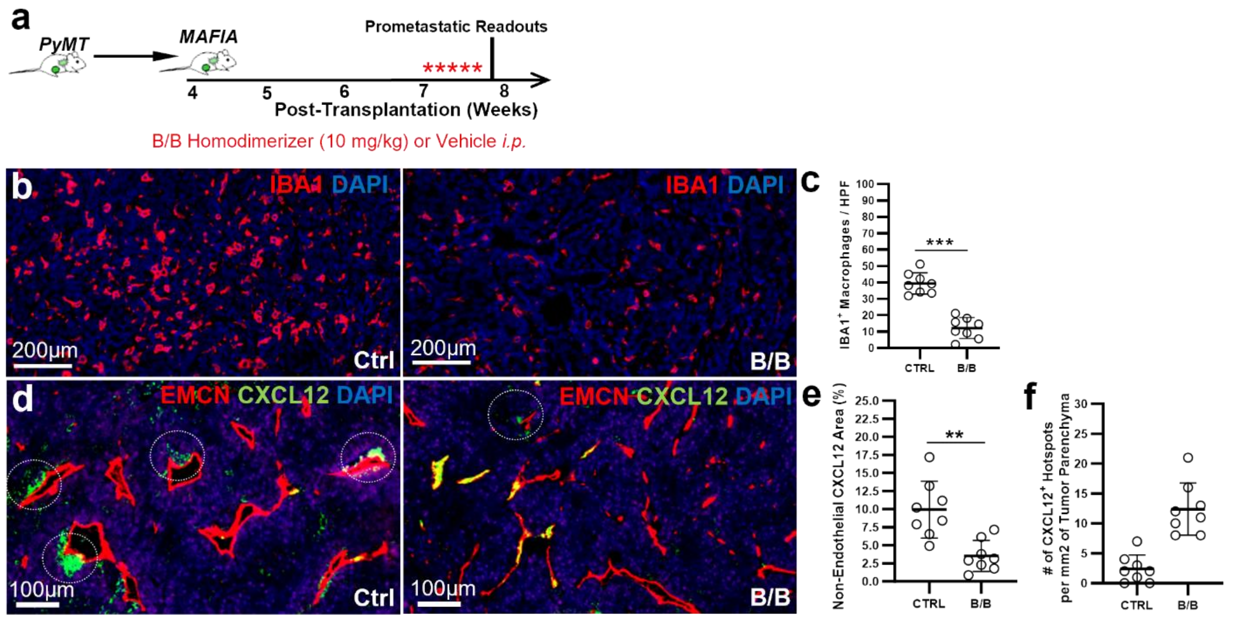


**Supplementary Figure 8. CXCL12 is partially derived from blood plasma. (a)** Experimental Strategy and pipeline for the development of MAFIA mice bearing PyMT tumor transplants. The B/B homodimerizer was given on a daily basis for 5 consecutive days to deplete macrophages, before sacrificing the animals. **(b)** Representative images of IBA1 immunofluorescence in the MAFIA-PyMT mice, to validate macrophage depletion. **(c)** Quantification of IBA1$^+$ macrophages via immunofluorescence in vehicle- and B/B-treated MAFIA-PyMT mice. *Mann-Whitney U-test.* **(d)** Representative images of CXCL12 expression using multichannel immunofluorescence. Note the CXCL12 "hotspots" at perivascular spaces, identified via green signal. However, when CXCL12 is not leaking through the vessel into the perivascular space and thus it overlaps with the EMCN$^+$ blood vessels, it can be identified via yellow signal. White circles reveal CXCL12 hotspots in perivascular spaces of Vehicle- and B/B-treated animals. **(e)** Quantification of CXCL12 area (%) in MAFIA-PyMT mice treated with either vehicle (Ctrl) or B/B, after excluding CXCL12 signal present within the EMCN$^+$ blood vasculature. *Mann-Whitney U-test.* **(f)** Quantification of the number of the CXCL12$^+$ hotspots identified in perivascular regions on the same fields-of-view, normalized per mm$^2$ of tumor parenchyma.

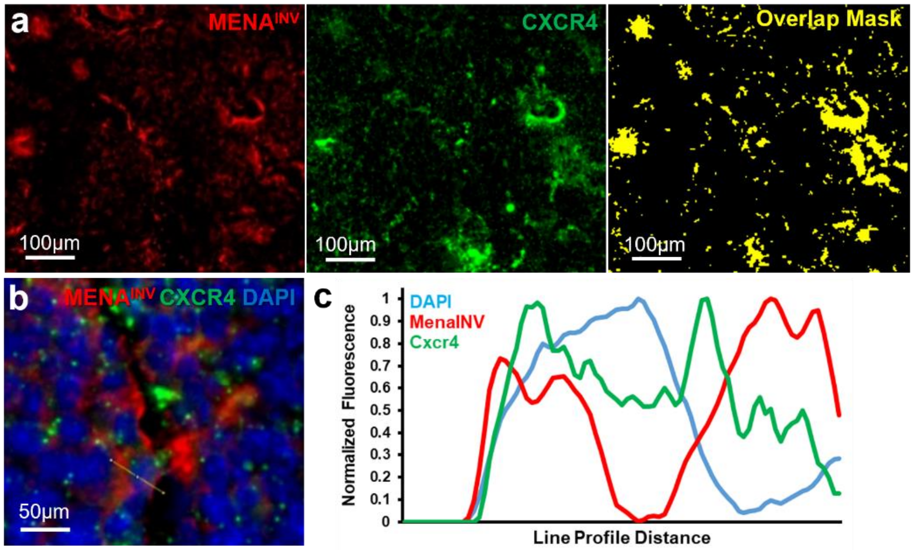


**Supplementary Figure 9. Co-expression analysis of CXCR4 and MENA$^{INV}$ in breast cancer.** **(a)** Multichannel immunofluorescence with CXCR4 and MENA$^{INV}$, revealing significant overlap between the two signals in the tumor microenvironment (shown as yellow "overlapping mask" in the third panel, after using an AND logic gate for MENA$^{INV+}$ and CXCR4$^{+}$ regions-of-interest; ROI). **(b-c)** Representative image at cellular resolution, showing Mena$^{INV+}$ tumor cells, also co-expressing CXCR4 (b). A line profile crosses through one such cell, and the relative intensities of the DAPI, MENAINV and CXCR4 signals are revealed along each pixel of the line (c). Note that the nuclear region (DAPI) has lower intensity of CXCR4 and MENA$^{INV}$ signals, while the highest intensity of these proteins is located on the edge of the cell, corresponding to plasma membrane and cytoplasmic region (c). All analyses of double-positive CXCR4$^{+}$MENA$^{INV+}$ cells has been verified through the line profiling approach. Analyses are shown here in the MMTV-PyMT mouse model.

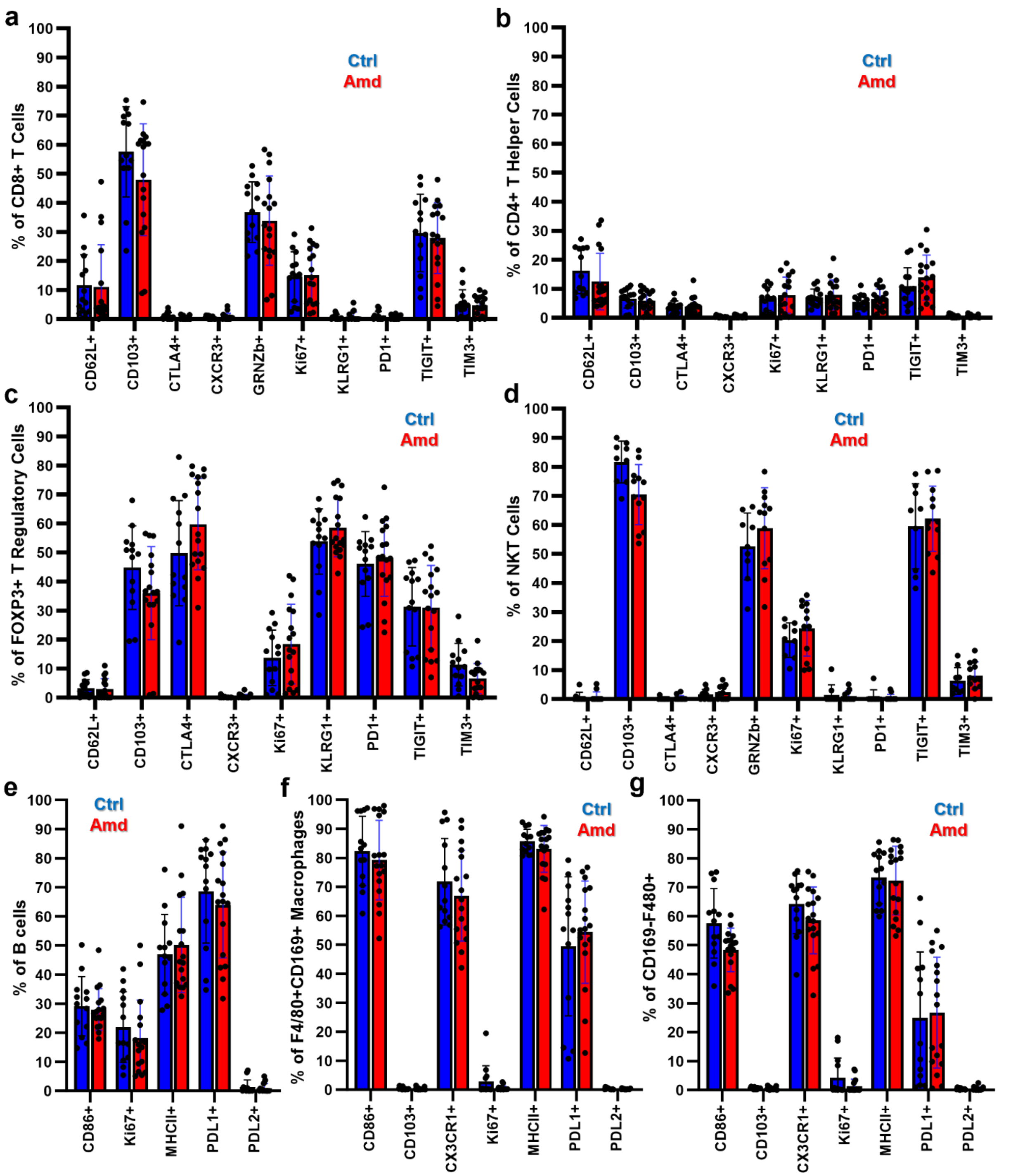


**Supplementary Figure 10. Expression of various phenotypic and functional markers in immune cell subsets using flow cytometry.** The immune cell subsets are defined based on gating strategy shown in Supplementary Figure 1. The immune cell populations that have been partially characterized in MMTV-PyMT mice receiving either the vehicle-control or Amd3100 are: **(a)** Cytotoxic $CD8^+$ T cells; **(b)** CD4+ T helper cells; **(c)** $FOXP3^+$ T regulatory cells; **(d)** NKT cells; **(e)** B cells; and **(f-g)** $F4/80^+CD169^+$ (f) or $CD169^-$ (g) macrophages. No changes are indicated in regard to the functional markers assessed after administration of Amd3100.

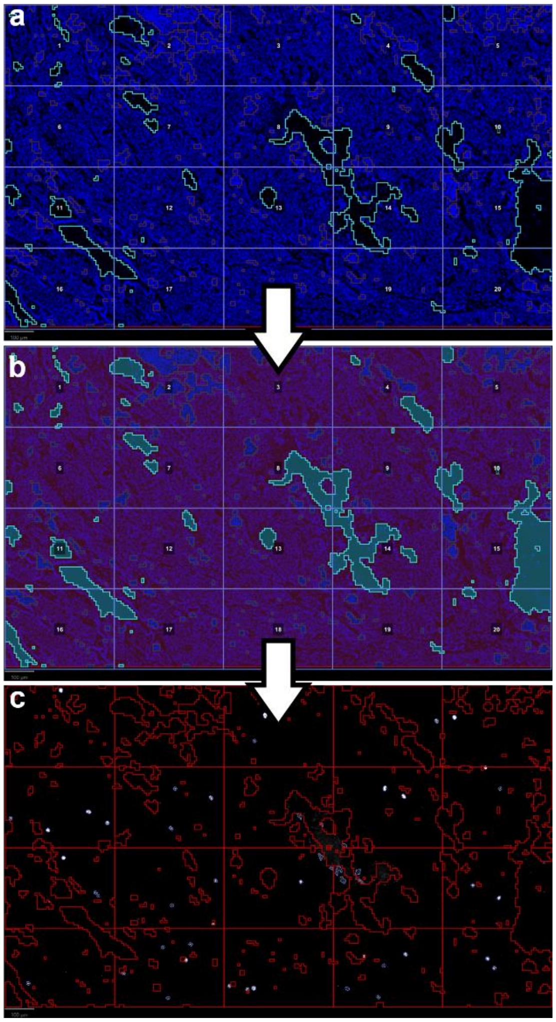


**Supplementary Figure 11. Pipeline for calculation of a homogeneity index, for CD8$^{+}$ T cell topology. (a)** Up to five representative 10x fields-of-view depicting the tumor area were captured. A “template ROI” is applied to the image, to segment the “DAPI ROI” into 20 equally-sized, rectangular-shaped sectors. **(b)** The “DAPI ROI” was automatically demarcated by a trained pixel classifier using some of the snapshots, in order to distinguish between DAPI cells (red areas), empty areas (cyan areas) and areas corresponding to dead cells/ necrosis (blue areas). **(c)** The CD8$^{+}$ T cell population was automatically demarcated and counted, using the cell detection tool. The density of the CD8$^{+}$ cells is then independently calculated for each sector. Subsequent mathematical approaches described in Materials and Methods utilize the densities of CD8$^{+}$ T cells in each sector to extrapolate the homogeneity of CD8$^{+}$ T cell distribution ($h_{CD8}$).
